# Trait Evolution in Microbial Communities

**DOI:** 10.1101/2020.12.15.422935

**Authors:** Lihong Zhao, Benjamin J Ridenhour, Christopher H Remien

**Affiliations:** Department of Mathematics and Statistical Science, University of Idaho, Moscow, ID, USA; Institute for Modeling Collaboration and Innovation, University of Idaho, Moscow, ID, USA

**Keywords:** Microbiome, trait dynamics, evolution, resource competition

## Abstract

Understanding the evolutionary dynamics of microbial communities is a key step towards the goal of predicting and manipulating microbiomes to promote beneficial states. While interactions within microbiomes and between microbes and their environment collectively determine the community composition and population dynamics, we are often concerned with traits or functions of a microbiome that link more directly to host health. To study how traits of a microbiome are impacted by eco-evolutionary dynamics, we recast a classic resource-mediated population dynamic model into a population genetic framework which incorporates traits. The relative fitness of each group of microbes can be explicitly written in terms of population dynamic parameters, and corresponding evolutionary dynamics emerge. Using several example systems, we demonstrate how natural selection, mutation, and shifts in the environment work together to produce changes in traits over time.

## 1 Introduction

Understanding the temporal dynamics of microbial communities is a key step towards the goal of predicting and manipulating microbiomes to promote beneficial states. It is critical to understand the evolution of microbial communities in response to perturbations such as from anitibiotics, diet shifts, and environmental changes. Such perturbations can have temporary and reversible effects or can permanently alter the microbiome. For example, the human gut microbiome can experience temporary reversible changes in species abundances without large gain or loss of bacterial species when a person travels between developed and developing countries, whereas enteric infection can lead to permanent decline and replacement of species [9]. On longer time scales, microbiomes may coevolve with their hosts [44].

While understanding how communities respond to perturbations is important for predicting dynamics, traits or functions of a microbiome often link more directly to health than community composition *per se.* Examples of such traits or functions include the protective effect against food allergies [6, 15], and the capacity to chemically modify ingested drugs [53]. The ability to model how key traits change with community composition is critical for engineering effective microbial communities that promote health or other beneficial states.

Evolution can affect microbial communities in several ways. Natural selection will drive adaptation to both the abiotic and biotic environment (which includes both the host and community structure). Changes in traits via natural selection have been extensively modeled, primarily via Price’s equation or its derivatives. The Price equation expresses the change in a trait as the covariance of the trait and fitness [38]. While natural selection erodes genetic variation around an optimum, mutation acts within the community to provide novel variants. This tension between selection and mutation should lead to mutation-selection balance in a community. Finally, evolutionary theory has shown the importance of genotype-by-environment (*G × E*) interactions in shaping traits [33, 13]. In the context of microbiomes, the community structure is a major contributor to the environment and can drive *G × E* effects. Changes in the abundances of species affects who interacts with whom and how much they interact, thus impacting coevolutionary dynamics via changing interaction strengths.

Resource competition models pioneered by MacArthur and Tilman have successfully been used to study macroecological communities [34, 50]. Indeed, many researchers are currently working to adapt these models for microbial systems. For example, Butler and O’Dwyer presented a consumerproducer-resource model for competitive interactions, and analyzed the local stability of equilibria of certain systems [7, 8]. Maslov et al. studied the assembly rules of microbial communities using conceptual models employing game theory methods [23, 24]. Their studies show that both the complexity and stability of microbial communities may arise from the mechanisms by which bacteria utilize resources.

In addition to competition and mutualistic cross-feeding, other types of interactions such as amensalism, commensalism, predation, and parasitism are important in microbiomes [1, 14, 36]. It has been repeatedly demonstrated that such interactions are context dependent and vary with the specific environment [4, 5, 11, 25, 32, 48]. This implies that the nature of interactions may vary over time and space. Thus, flexible modeling frameworks—such as Tilman’s [50]—which can encompass numerous types of interactions are ideal.

While modeling studies have undoubtedly advanced our understanding of microbial community dynamics, trait dynamics have been largely unexplored. In this paper, we recast a classic resource-mediated dynamic model into an evolutionary framework that relies on the Price equation [38]. This framework of connecting population dynamic models to evolution has been successfully applied in epidemiology to study, for example, the evolution of virulence [10, 18, 19]. We group microbes by their functional characteristics, and use the Malthusian fitness of each strain to model the eco-evolutionary dynamics within the community. Transforming the system in this way reveals how model parameters affect the three evolutionary forces driving dynamics: natural selection, mutation, and changes in the environment. Incorporating traits into this framework yields a form of the Price equation that tracks the dynamics of the mean value of any trait of interest. We begin by introducing a general resource-mediated model in Section 2. Section 3 illustrates the population-genetic and trait-based approach. This modeling framework allows us to make predictions about trait evolution, and to model the non-equilibrium ecological dynamics.

## 2 Mathematical Model

We extend Tilman’s model of resource competition [50] to incorporate other types of interaction and the production of metabolic byproducts which may also mediate population dynamics.

### 2.1 General model

For a community of *M* groups of microbes with abundances 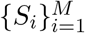 and *N* resources with abundances 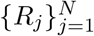, the general model is given by

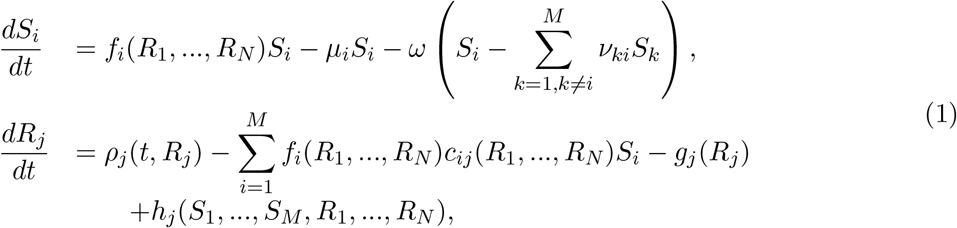

for *i* = 1, …,*M* and *j* = 1, …,*N* (see Table 1 for main notations). Note that the identity of each group of microbes in this model is determined by its dependence of per capita reproductive rate on available resources *f_i_*(*R*_1_,…, *R_N_*) and mortality rate *μ_i_*. Our model defines {1,2,…,*M*} as a set of biological ‘groups’; groups could, for example, be a set of strains of one species, a set of different species, or a set of *m* species (*m* < *M*) with some species having more than one strain. We can impose conditions such that inter-species mutations are not allowed. Assume M groups of microbes consist of *L* species *(L* < *M*), let *U_i_* be the collection of groups/strains of species *i*, and 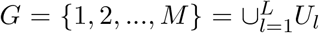 with 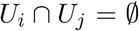 for *i* ≠ *j*. If microbes in group i are of one strain of species *I, i ∈ U_I_*, then *ν_ki_* = 0 for *k* ∈ *G* \ *U_I_* and ∑_*i≠k*_ *ν_ki_* = 1.

**Table 1:** Main notations used in model (1).

| Notation | Description |
| --- | --- |
| *Variables* |  |
| $R_j$ | Abundance of resource $j$ |
| $S_{ij}$ | Abundance of microbial group $i$ that consumes resource $j$ |
| $S_i = \sum_{k=1}^N S_{ik}$ | Abundance of microbial group $i$ |
| $S = \sum_{l=1}^M S_l$ | Total microbial population abundance |
| $p_{ij} = \frac{S_{ij}}{S_i}$ | Proportion of microbes in group $i$ that consumes resource $j$ |
| $q_i = \frac{S_i}{S}$ | Proportion of microbial group $i$ within total population |
| $\eta_{ij}$ | Instantaneous per capita rate of change of the abundance of $S_{ij}$ in the absence of mutation/Malthusian fitness of $S_{ij}$ |
| $\bar{\eta}_i = \sum_{k=1}^N p_{ik} \eta_{ik}$ | Average instantaneous per capita rate of change of the abundance of $S_i$ in the absence of mutation/Malthusian fitness of $S_i$ |
| $\bar{\eta} = \sum_{l=1}^M q_l \bar{\eta}_l$ | Average instantaneous per capita rate of change of the abundance of $S$ in the absence of mutation/Malthusian fitness of $S$ |
| $\Delta_{ns}$ | Effect of natural selection on the change in $\bar{\eta}$ |
| $\Delta_m$ | Effect of mutation on the change in $\bar{\eta}$ |
| $\Delta_{ce}$ | Effect of the change in the environment on the change in $\bar{\eta}$ |
| *Functions* |  |
| $f_i(R_1, \dots, R_N)$ | Dependence of the per capita reproductive rate of group $i$ on resource availability |
| $\rho_j(t, R_j)$ | External influx rate of resource $j$ |
| $c_{ij}(R_1, \dots, R_N)$ | The amount of resource $j$ required to produce each new individual of group $i$ |
| $g_j(R_j)$ | Depletion rate of resource $j$ due to degradation and/or outflow |
| $h_j(S_1, \dots, S_M, R_1, \dots, R_N)$ | Export rate of resource $j$ as a metabolic byproduct |
| *Parameters* |  |
| $\omega$ | Mutation rate |
| $\nu_{ki}$ | probability that $S_k$ mutate into $S_i$ |
| $\mu_i$ | Mortality rate of microbes in group $i$ |
| $\rho_j$ | Influx rate of resource $j$ |
| $\sigma_j$ | Degradation rate of resource $j$ |
| $\alpha_{ij}$ | Uptake rate of resource $j$ by microbes in group $i$ |
| $\epsilon_{ij}$ | Conversion efficiency of resource $j$ by microbes in group $i$ |
| $\beta$ | Resource switching rate |
| $\gamma_i = \epsilon_i \alpha_i$ | Net per capita reproductive rate of microbes in group $i$ |

Let 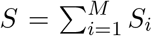 denote the total abundance of the microbial population. The change in the proportion of group 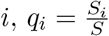, can be tracked with

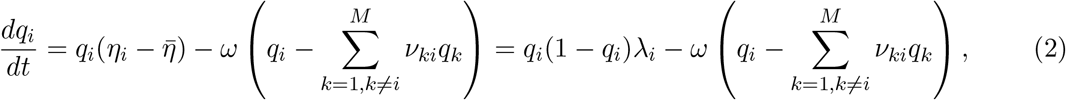

where

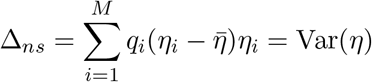

is the instantaneous per capita rate of change of the abundance of group i in the absence of mutation (Malthusian fitness of *S_i_*), 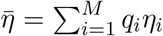 is the average Malthusian fitness of *S*, 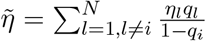 is the average Malthusian fitness of *S* — *S_i_*, and 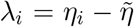 is the selection coefficient of group *i*. Neglecting mutations, from equation (2) we can see that when there is at least one group present along with group *i*, the frequency of group *i* will increase (or decrease) when the fitness of group *i* is higher (or lower) than the average fitness of all other groups.

With equation (2), we can derive the following equation for the temporal dynamics of 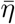:

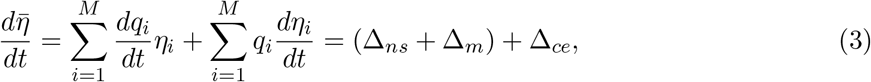

where

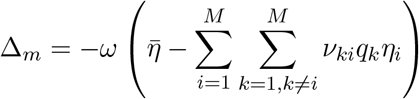

denotes the effect of natural selection on the change in 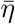. The term Δ_*ns*_ is equal to the variance in *η* across all groups and is always non-negative. The term

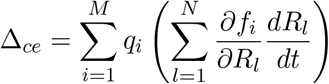

denotes the effect of mutation on the change in 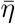. It is negative if the average fitness of the mutants 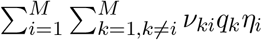 is lower than the average fitness of the total population 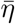. The term

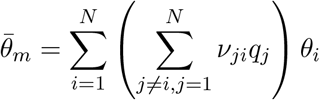

denotes the effect of the change in the environment on the change in 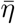. Note that the *environment* includes all the forces (abiotic and biotic factors) other than the force of natural selection at the specific level. Natural selection tends to increase the mean fitness 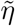, but mutation and change in the environment may have negative impact on the mean fitness. These effects will balance at equilibrium.

### 2.2 Trait Dynamics

In addition to the dynamics of the frequencies of different groups of microbes and the dynamics of mean fitness of the population, we are also interested in the dynamics of microbial phenotypic characteristics that may impact the ability of microbes to survive in a specific environment. These characteristics can be morphological, physiological, or behavioral traits such as shape or color of bacterial colonies, maximal growth rate, the ability to metabolize carbon compounds, or the ability to survive at different pH level. Let *θ* denote a quantitative trait of interest. Each group within the microbiome has a specific value of the focal trait at time *t, θ_i_*(*t*), and the mean trait value across the microbiome, 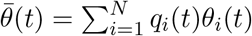, which is affected by the population dynamics of the microbiome. We can track the dynamics of the mean trait value 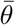 with

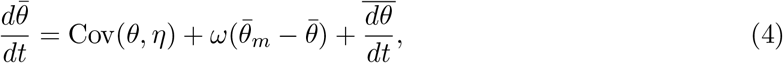

where

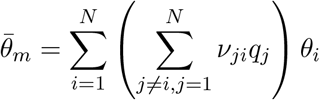

is the average value of trait *θ* among all new mutants, and

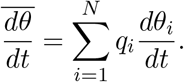

Equation (4) is a continuous-time derivation of Price’s equation [38] which is simple but informative. It tells us that the change in the average value of trait *θ* is driven by three processes given by the three terms in Eq. (4). First, natural selection on *θ* is given by the covariance between the trait and fitness across all groups. The second term is change due to mutation which scales with the mutation rate and is positive if the average trait value among mutants is larger than the average trait value of the total population at a given time. The third term represents other factors that affect the trait value of each strain (e.g., drift, environmental effects).

### 2.3 Alternative Form and Multi-level Selection

In microbiome studies, it is common to group organisms at different taxonomic levels. Often operational taxonomic unit (OTU)-based classification is at family, genus, or species levels. It has been observed in microbiology that strains of a single species vary in their functional capacity, such as drug resistance, virulence, or ability to uptake different compounds from environment [26, 40]. These differences among strains may also impact the health of hosts [12]. Motivated by this, we can rewrite the model in Section 2.1 for the case that individuals in group *i* can be further divided into subgroups *ik*, such that *k* = 1, …,*L_i_*. Here, the group and related subgroup could be, for example, genus and species or species and strain.

Let *S_i_* denote the abundance of group *i* and *S_ik_* denote the abundance of subgroup *ik*, with 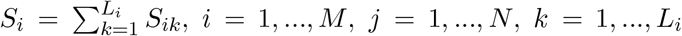, where *L_i_* denotes the total number in subgroups of group *i*. Define *p_ik_* = *S_ik_/S_i_* as the frequency of subgroup *ik* within group i, and *q_i_ = S_i_/S* as the frequency of group *i*. Denote 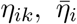, and 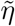 as the Malthusian fitness of *S_ik_, S_i_*, and *S*, respectively. The temporal dynamics of 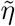 can be written as

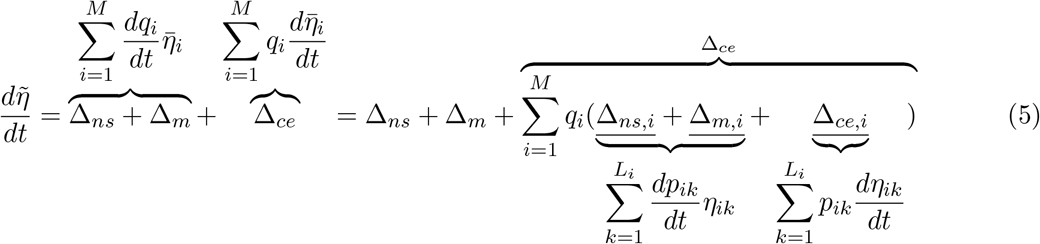

(derivation in Appendix A). Similar to equation (3), the change in 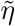 is driven by three processes: natural selection between groups (Δ_*ns*_), mutation (Δ_*m*_), and change in the environment (Δ_*ce*_). The effect of change in the environment can be further decomposed into the same three processes:

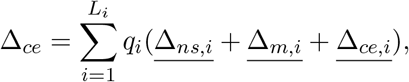

where 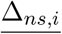 is natural selection between subgroups, 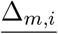 is mutation, and 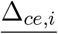) is the change in the environment, for each group *i*.

In this case, the dynamics of 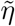 are driven by selection at two levels: selection between groups (Δ_*ns*_, Δ_*m*_, Δ_*ce*_) and selection between subgroups 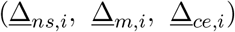. Equation (5) can be expanded recursively to represent the change in the mean fitness at different levels of nested groups, which provides a hierarchical decomposition of selection within and between groups.

## 3 Simulation Studies

Some in vitro studies suggest that negative interactions dominate synthetic aquatic microcosms [16] and human microbiota [51]. These negative interactions can be the result of competition for resources (e.g., nutrients, space) or damage caused by toxins [21, 28, 40]. Some studies suggest that cooperation and higher-order interactions also impact the functioning of microbial communities [3, 39].

In this section, we apply the approach developed in Section 2 to several models to illustrate the advantages of this approach. We present simple communities in which microbes compete for abiotic resources without the production of metabolic byproducts. Cases with metabolic byproducts are found in Appendix C. These simple communities highlight how interactions affect selective pressures over time.

### 3.1 Competition for a single resource

Microbes can compete for resources such as nutrients, light, or space. These competitive interactions can occur between a broad genetic range of microbes, from similar strains to members of different phyla [40]. As a simple example, consider a community of *N* groups of microbes with abundances 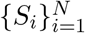 competing for one externally supplied resource *R*:

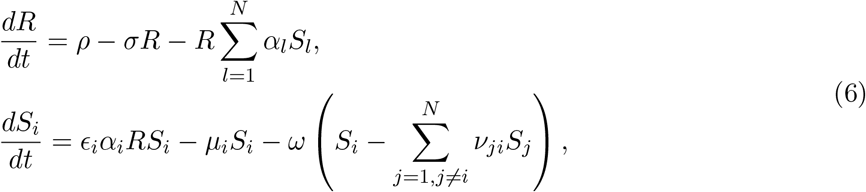

for *i* = 1,…, *N*. Here, we assume a linear functional response for growth rate and *σ* = 0. In this case, *η_i_* = *ϵ_i_α_i_R* – *μ_i_*, and 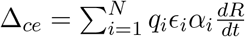.

Note that in our model, the fitness of microbes in a given group depends on the net per capita reproductive rate *γ_i_* = *ϵ_i_α_i_*, mortality rate *μ_i_*, and resource level *R*, with the parameters *γ_i_* and *μ_i_* driving the evolutionary dynamics of the community. Reproductive and mortality rates of a given group can be thought of as traits. Utilizing equation (4), we can track the dynamics of 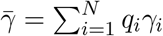 and 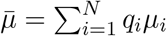 with

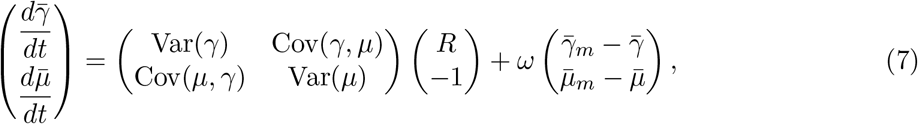

where 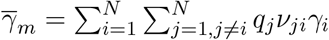 and 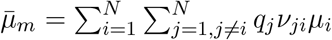 are the average net per capita reproductive rate and mortality rate among all the mutants, respectively, and (*R*, — 1)^*T*^ is the selection gradient. This equation is analogous to quantitative genetics models [29]. It shows that natural selection favors increased net per capita reproductive rates and the strength of selection is proportional to the resource abundance, *R*. It also shows that reduced mortality rate is favored with a selection strength of −1. The evolution of each trait is constrained by the genetic variance in the focal trait (direct selection), possible covariance between the focal trait and other traits (indirect selection), and mutation. It is important to note that the direction of natural selection changes with changes in the selection gradient (*R*, – 1)^*T*^ (i.e., *R* changes in magnitude over time). Thus, the dynamics of resource and microbial population abundances are driven by both ecological and evolutionary processes.

When mutation is absent, at equilibrium, model (6) gives 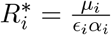 with one value of 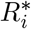 for each group *i*. Theory predicts that the group with the lowest value of 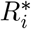 will competitively displace all other groups at equilibrium. Two or more groups coexist at equilibrium only if they have identical value of 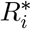 [49, 50]. More generally, n groups cannot coexist on fewer than n resources—the ‘competitive exclusion principle’ [2].

Traditional equilibrium and near-equilibrium analyses focus on predicting properties of equilibria. The approach we introduced in Section 2 allows us to predict the transient evolutionary dynamics, which complements the insights gained from equilibrium and near-equilibrium analyses.

#### Example 1

The first example is a community of three groups of microbes competing for one single resource; key parameter values and initial conditions are listed in Table 2. When mutation is absent, *S*_3_ outcompetes *S*_1_ and *S*_2_ because it has the highest fitness and consequently the lowest value of 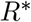. When mutation is present (Δ_*m*_ ≈ 0), all three groups of microbes co-exist with *S*_1_ and *S*_2_ maintaining low abundances. Different evolutionary forces dominate at different stages, and at steady state these forces balance. In particular, when mutation is present, mutation-selection balance is reached and selection is nonzero at equilibrium.

**Table 2:** Parameter values and initial conditions used in Figure 1.

| Parameter | Value | Parameter | Value | Parameter | Value | Parameter | Value |
| --- | --- | --- | --- | --- | --- | --- | --- |
| $\rho$ | 3.6 | $\mu_1$ | 0.7 | $\mu_2$ | 0.7 | $\mu_3$ | 0.8 |
| $\{\epsilon_i\}_{i=1}^3$ | 0.6 | $\alpha_1$ | 0.6 | $\alpha_2$ | 0.4 | $\alpha_3$ | 0.8 |
| $\nu_{12}$ | 0.5 | $\nu_{13}$ | 0.5 | $\nu_{21}$ | 0.7 | $\nu_{23}$ | 0.3 |
| $\nu_{31}$ | 0.4 | $\nu_{32}$ | 0.6 | $R(0)$ | 2 | $\{S_i(0)\}_{i=1}^3$ | 1 |

We also track the dynamics of three different traits: net per capita reproduction rate *γ* = *ϵα*, mortality rate *μ*, and a trait *θ* that is not directly associated with the fitness of each strain (*θ*_1_ = 0.9, *θ*_2_ = 0.3, and *θ*_3_ = 0.3) (Figure 2). Eq. (4) shows that the dynamics of a trait depend critically on the covariance of the trait and fitness across all groups. At equilibrium, mutation and natural selection will balance. The trait *θ* is not directly associated with the fitness of microbes. For these parameter values, the group with smallest value of *θ* has the highest fitness so that Cov(*θ, η*) < 0. Natural selection favors *S*_3_ which has lowest value of *θ_i_*, leading to a decrease in 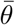. In this case, the average trait value among all new mutants 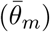 is higher than the average trait value of the entire population 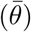, leading to an increase in 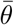. Natural selection is the dominant evolutionary force acting on *θ*.

**Figure 1:**
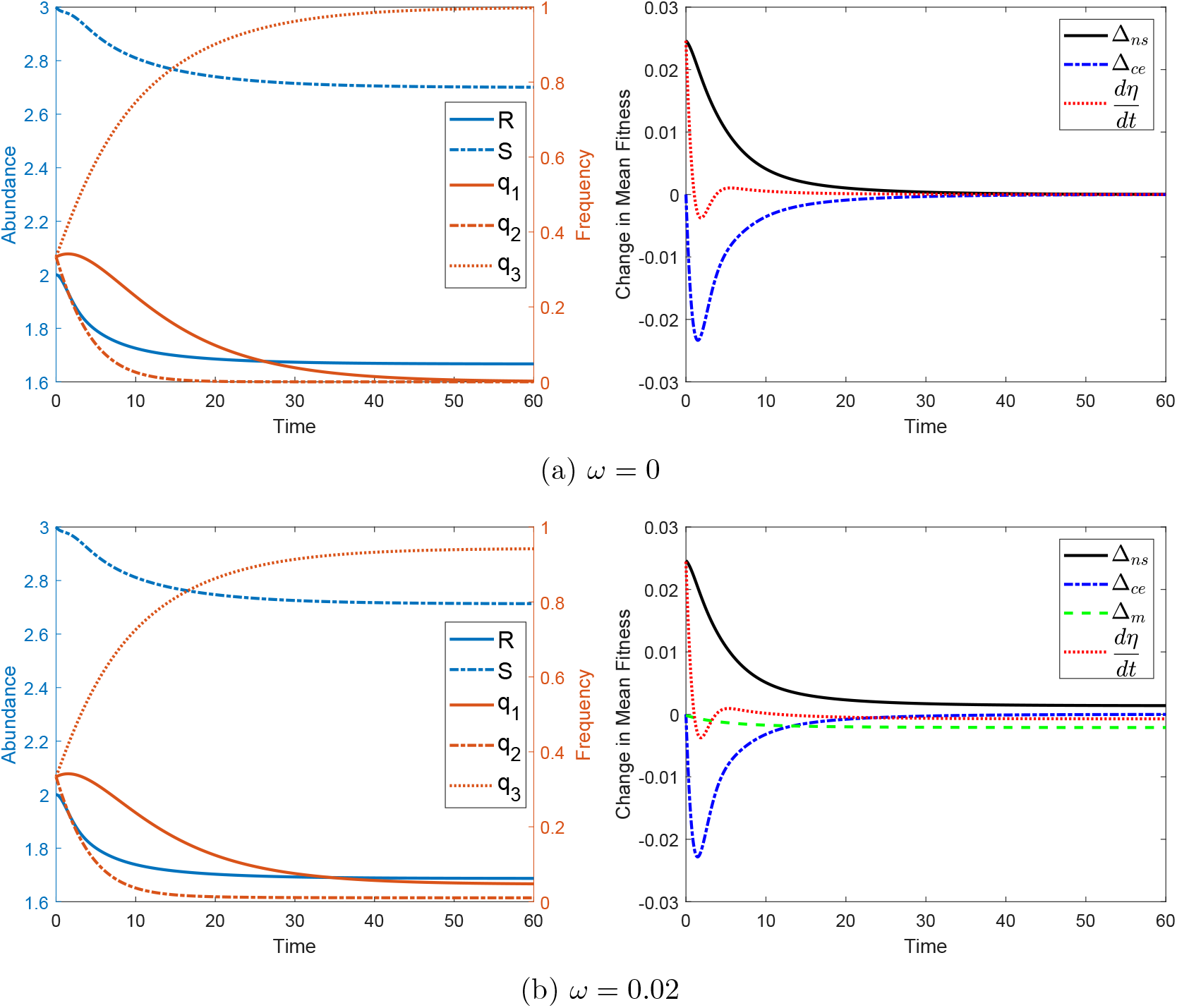
Resource competition without and with mutation. Numerical simulation of Example 1. In (a), there is no mutation, while in (b), we allow microbes to emerge by mutation (*ω* = 0.02). From (a) to (b), we show the effect of mutation. The left panels show the abundance of resources and total microbial population (left axis) and frequencies of each group of microbes (right axis) over time, the right panels show the change in mean fitness 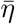 (red dotted line) and effects of different drivers over time (black solid line: Δ_*ns*_, blue dotted line: Δ_*ce*_, green dash line: Δ_*m*_).

**Figure 2:**
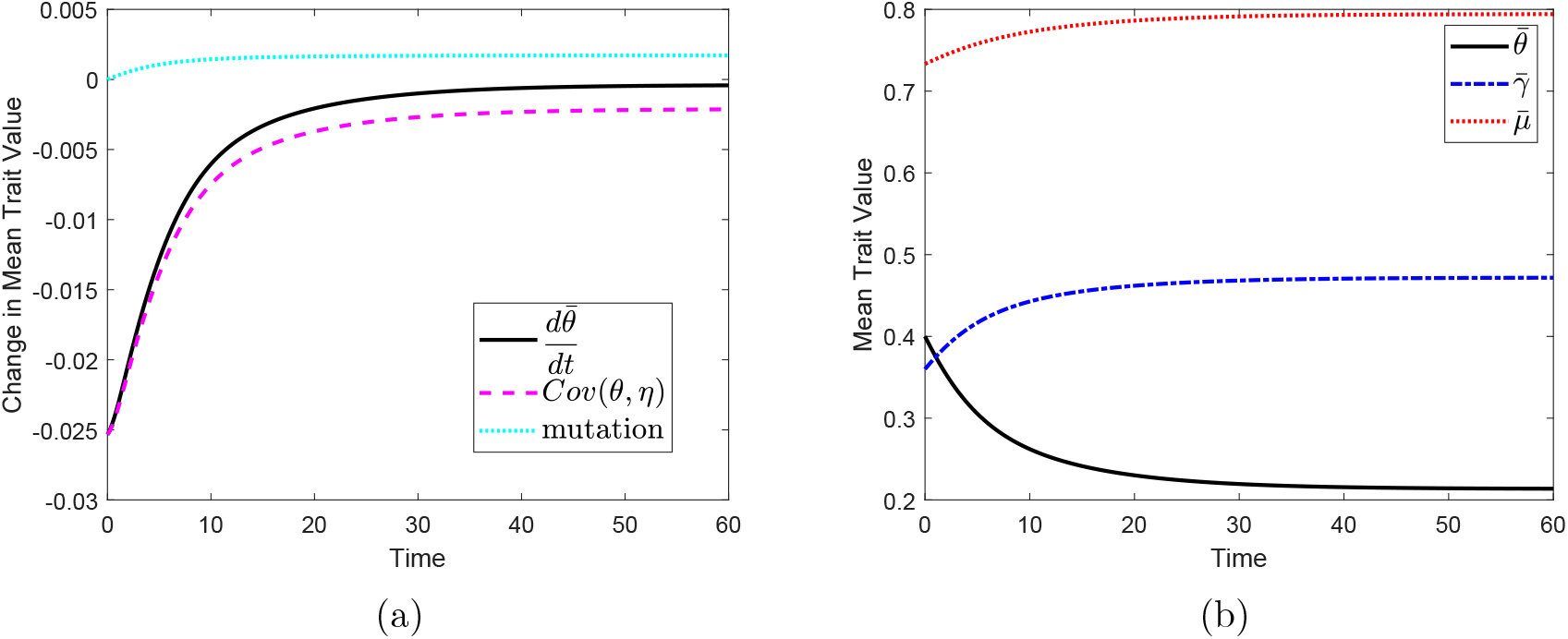
Resource competition with trait evolution, Example 1 with mutation (the community shown in Figure 1b). (a) Rate of change in the average value of a focal trait *θ* (black solid line: 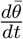), and effect of selection govern by different forces (cyan dotted line: mutation, magnate dash line: natural selection Cov(*θ, η*)). (b) Mean value of different traits over time: net per capita reproductive rate *γ = ϵα* (blue dotted-dash line), mortality rate *μ* (red dotted line), and trait *θ* (black solid line).

As for the other two traits that are directly associated with microbial fitness, large values of *γ* and small values of *μ* lead to high fitness. For both of these two traits, effects of natural selection are positive and effects of mutation are negative, with natural selection being the dominant force driving trait evolution initially, while mutation is dominant later.

### 3.2 Resource competition with variable resource supply rate

Normally microbes do not grow in an environment where resources are supplied at a constant rates. We consider two types of variations in resource availability, the first imposes a temporary shift and and the second continuous oscillations. In reality, the resource supply rate could vary in any number of ways, but these two types of variations provide insight into how two types of perturbations affect dynamics. For simplicity, we use the community shown in Figure 1b as the baseline, start the simulation with initial condition (*R*(0), *S*_1_(0), *S*_2_(0), *S*_3_(0)) = (1.687, 0.126, 0.028, 2.559), close to the equilibrium in the baseline system, and then perturb the system.

#### 3.2.1 Temporary perturbation

First, we perturb the resource supply rate to incorporate a temporary decrease:

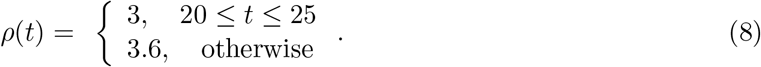

The temporary decrease in external resource supply rate leads to significant changes in resource and microbial abundances that deviate from the original steady state. At the later phase of this temporary decrease in resource supply rate, the rate of decrease in microbial abundance slows down as resource starts to build up, and the system approaches a new steady state towards the end of this period. As the resource supply goes back to the original rate, microbial abundance starts to increase once resource abundance is sufficiently high, and the system eventually returns to the original equilibrium state as in Figure 1. During and after the perturbation, environmental change is the dominate force driving the dynamics of mean fitness, followed by natural selection, as shown in Figure 3a. Note that in this case, perturbation has little effect on the proportions of different groups of microbes. As we assume that the system is well-mixed, individual microbes have equal access to resource, thus little fluctuation in mean trait value.

**Figure 3:**
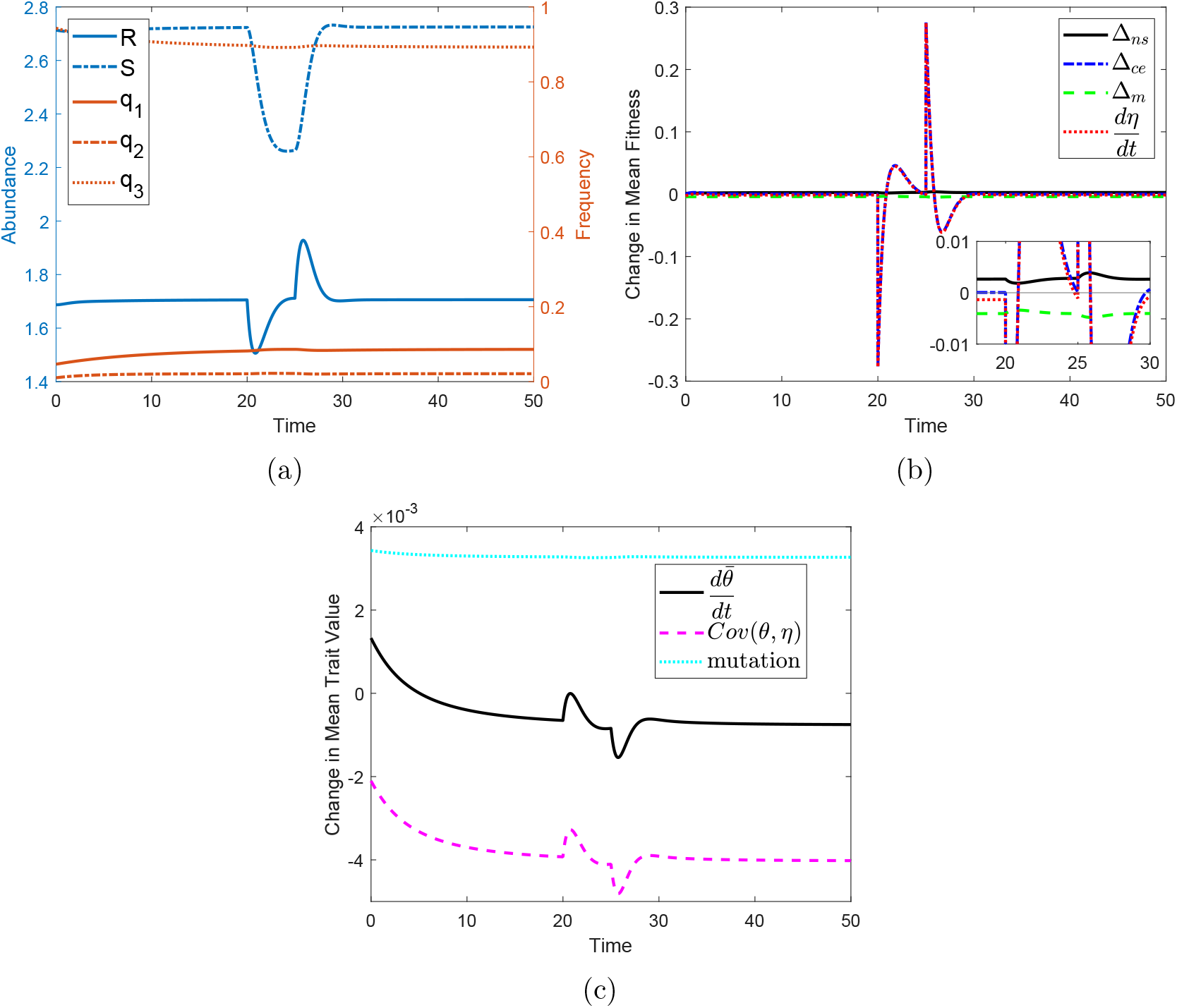
Numerical simulation of Example 1 with **non-constant resource supply rate given in** (8), initial condition is (*R*(0), *S*_1_(0), *S*_2_(0), *S*_3_(0)) = (1.687, 0.126, 0.028, 2.559), other parameter values are the same as used for Figure 1b.

#### 3.2.2 Oscillatory resource supply rate

As shown in Figure 4, oscillatory resource supply rate 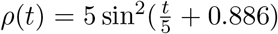 leads to oscillatory behaviors in abundances, mean trait values, and different evolutionary forces. The change in mean fitness is mainly driven by changes in environment, and the change in mean trait value is mainly driven by selection as the microbial fitness depends on resource availability.

**Figure 4:**
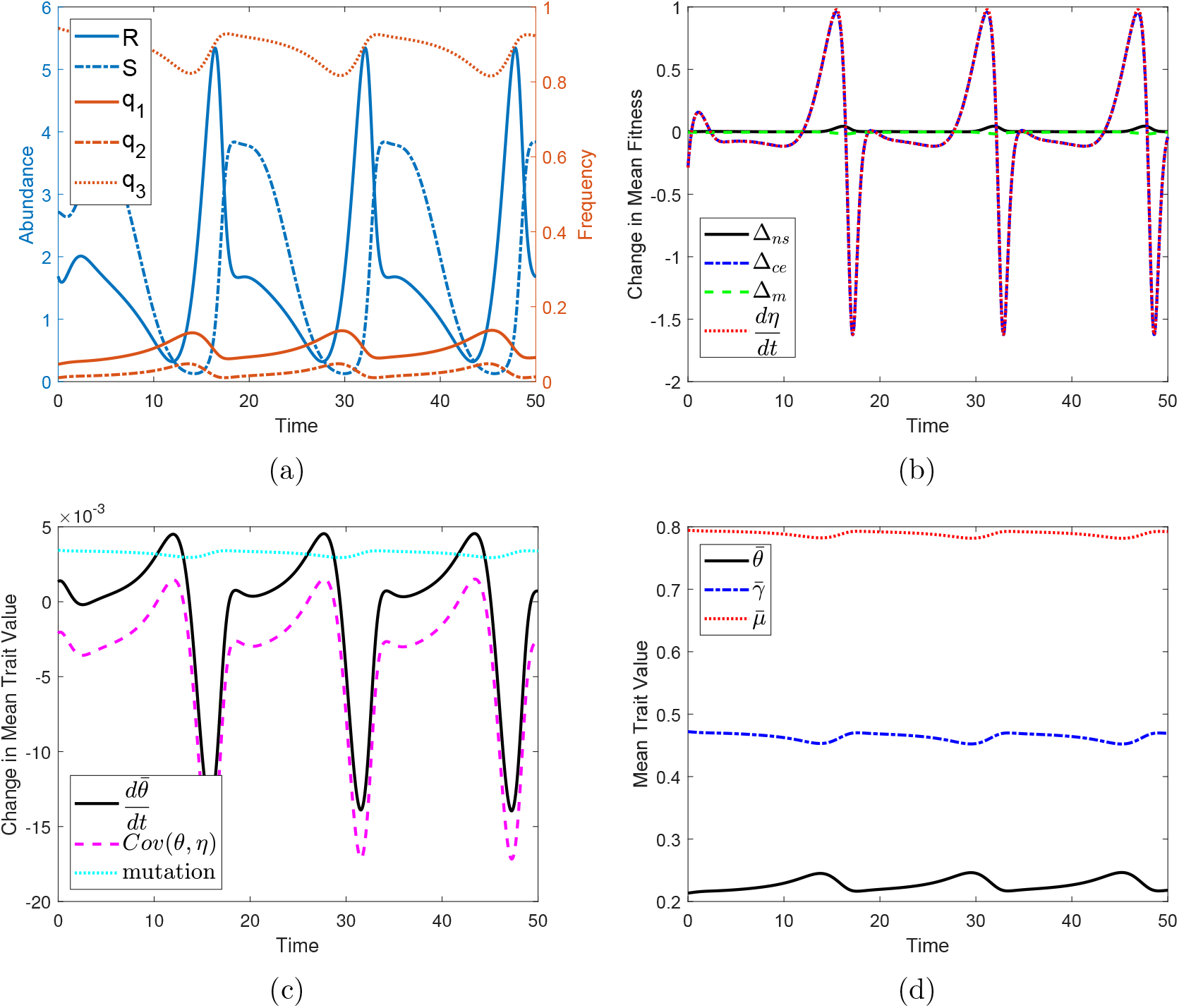
Numerical simulation of Example 1 with **periodic resource supply rate** 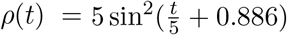, initial condition is (*R*(0), *S*_1_(0), *S*_2_(0), *S*_3_(0)) = (1.687, 0.126, 0.028, 2.559), other parameter values are the same as used for Figure 1b.

### 3.3 Resource switching

Carbon is one of the main resources for bacterial growth [45] but may come in many different forms (e.g., there are many types of sugars). When multiple carbon sources are present, bacteria display two types of growth behavior: these carbon sources can either be simultaneously consumed (e.g., co-utilization or co-metabolization) or be utilized in a hierarchical manner (e.g., carbon catabolite repression) [22]; in other words, bacteria demonstrate preferences for particular carbon sources. Likewise, bacteria can also switch between aerobic and anaerobic metabolic pathways which have different resource utilization [52] but typically prefer aerobic metabolism when feasible.

As a toy example of metabolic mechanism switching, we construct a community of *M* groups of microbes and *N* interchangeable resources such that each group *i* corresponds to a distinct species, assume any individual can only consume one resource at a time, and individuals are allowed to switch between different resources based on their preference and resource availability. We can track the abundance of species *i* consuming resource *j* (i.e., strain *j* of species *i*), *S_ij_*, and the abundance of resource *j, R_j_*, using the following model:

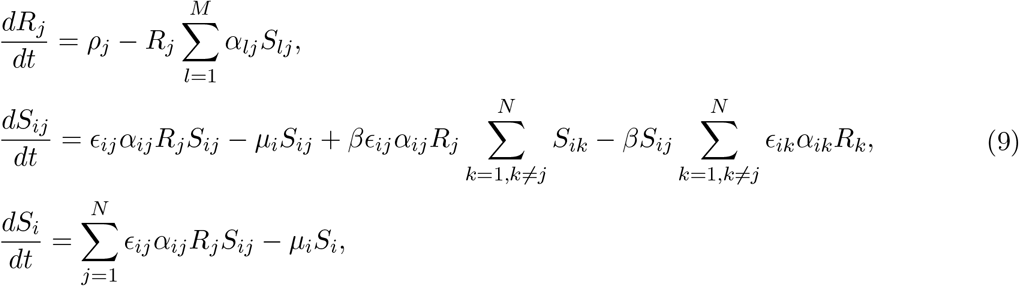

for *i* = 1,…, *M*, and *j* = 1,…, *N*, where *ρ_j_* is influx rate of resource *j, α_ij_* represents the rate of consumption of resource *j* by species *i, ϵ_ij_* characterizes the resource conversion rate of species *i* on resource *j*, and *β* represents the resource switching rate.

Similar as in Section 2.3, let 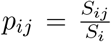 be the frequency of strain *j* of species *i*, 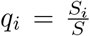 the proportion of species *i* to the total population, we can rewrite the model (9) as

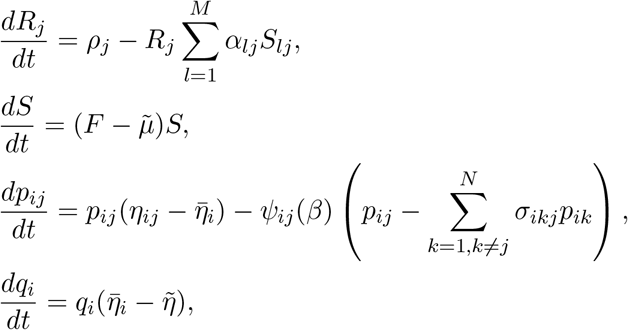

where 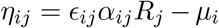 is the Malthusian fitness of *S_i_*; 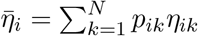 is the mean Malthusian fitness of 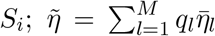 is the mean Malthusian fitness of *S*, 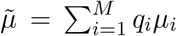 is the average mortality rate over the entire population, 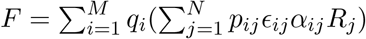 is the average per capita reproduction rate over the entire population,

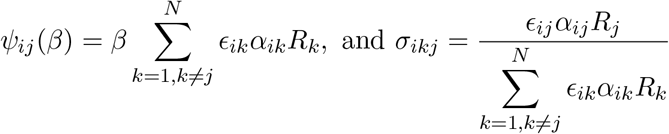

are analogous to the mutation rate, 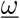, and probability that *S_il_* mutate into 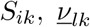, in model (10), respectively. Note that *ψ_ij_* (*β*) and *σ_ikj_* are time-dependent here. Similarly, differentiating the expression for 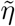 with respect to time, we obtain

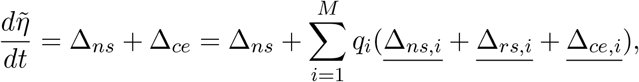

where

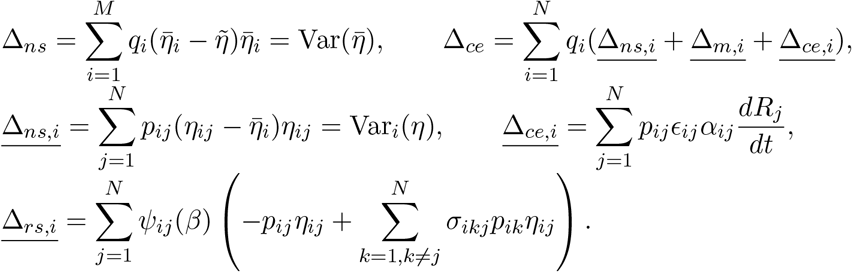

Note that 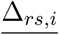 here denotes the effect of resource switching on the change in 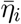, and it is analogous to 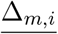 in equation (5). If microbes are not allowed to switch to a different resource, *β* = 0, then *ω*(*β*) = 0 and 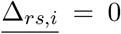, the temporal dynamics of 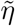 is driven by natural selection and changes in the environment only. Further, if we assume the dynamics of resources are much faster than the dynamics of microbial population and apply separation of timescale, i.e., set 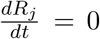 for *j* ∈ {1, …,*N*} then substitute the resulting algebraic constraints into 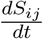, the environmental dependence of 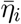 is absent, 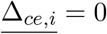, and the environmental dependence of 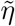 is simplified into the effect of natural selection between subgroups, 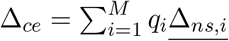.

#### Example 2

Consider a community of 3 microbial species and 2 substitutable resources, in which individuals can only consume one resource at a time and can switch between those two resources based on their preference, as shown in Figure 5. For simplicity, we assume that the resource conversion rates are the same for all the species on all resources, *e_ij_* = *ϵ* for all *i* and *j*, with *M* = 3 and *N* = 2 in model (9).

**Figure 5:**
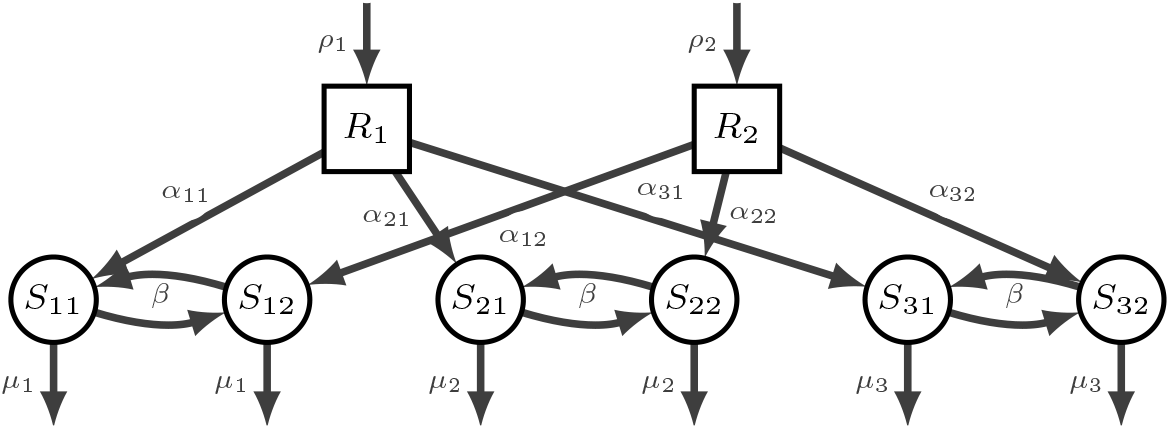
Illustration of the topological structure of the microbial community discussed in Example 3.

When resources are supplied at constant rates and resources are abundant, the community reaches equilibrium state, decreasing resource supply rate will increase the time it takes for the community to reach equilibrium, if there is one, and can lead to instability when the resource supply rate is sufficiently low (simulation results not shown here). If the supply rate of one resource is time-dependent, *ρ*_1_(*t*) for *R*_1_ in this case, the abundances of the other resource and all microbial species as well as strain frequencies also isolate over time, as shown in Figure 13 (note that *R*_2_ appears to reach equilibrium state in Figure 13a, but it actually fluctuate over time with much smaller magnitude). *S*_2_ and *S*_3_ dominate the population at different time intervals while *S*_1_ remains at low abundance and eventually approaching zero. From Figure 6 we can see that the change in mean fitness for different species fluctuate as well due to the time-dependent supply of *R*_1_, and slight time delay is observed in the effects of natural selection and mutation in comparison to the change in the environment. The magnitude of changes in individual species fitness are different, the magnitude of changes in 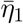 is the highest, and 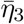 is the lowest. If the system begins away from equilibrium state, the patterns of evolutionary dynamics between different species are different (results not shown here).

**Figure 6:**
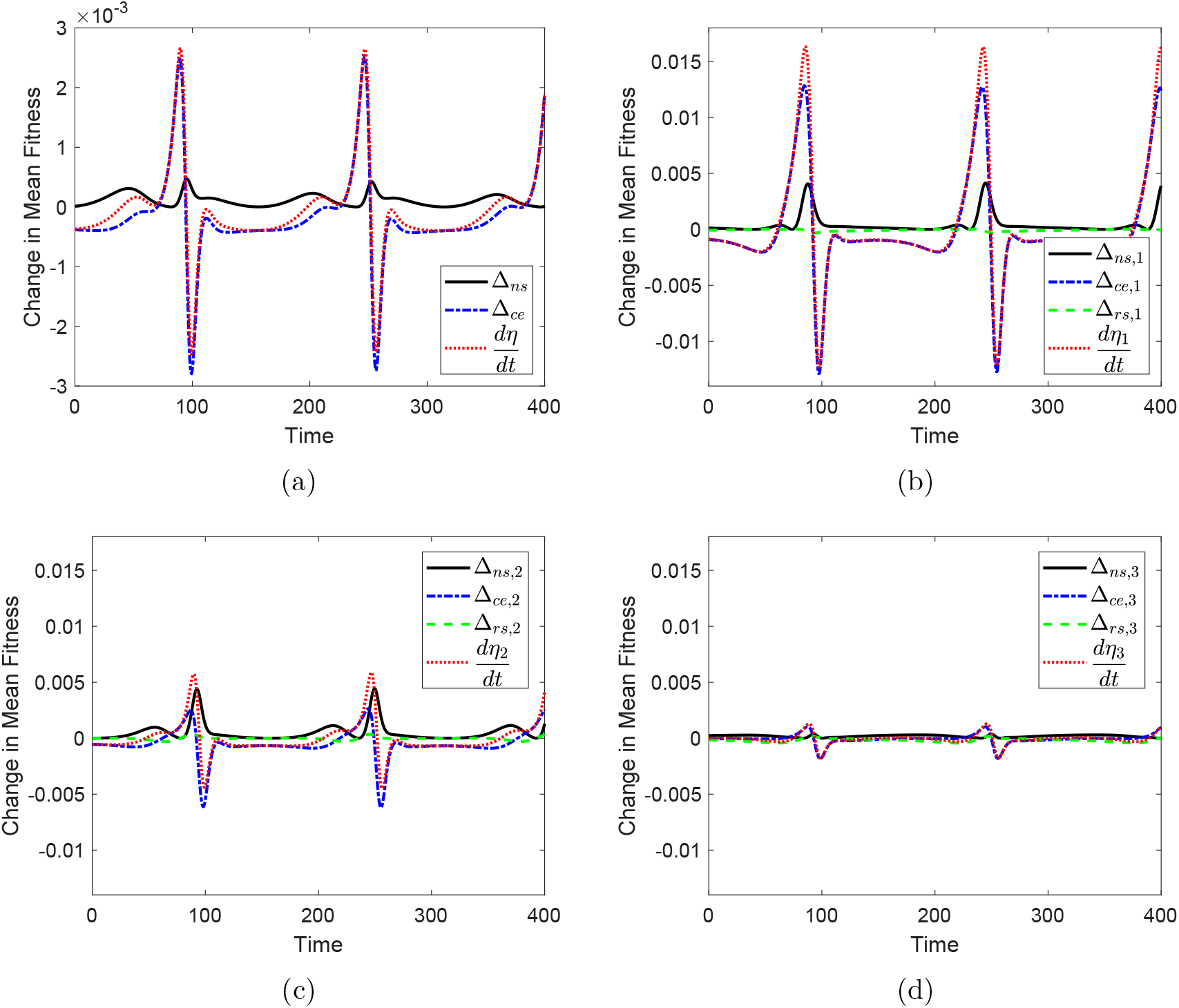
Numerical simulation of the community in Example 3. The evolutionary forces on mean fitness can be decomposed at both the species level ((a)) and strain level ((b)-(d)). In (a) we plot the change in mean microbial population fitness, 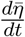 (red dotted lines), and distinguish the effects of natural selection, Δ_*ns*_ (black solid line), and changes in the environment, Δ_*ce*_ (blue dotted-dash line). In (b)-(d), we plot the changes in mean fitness of each group of microbes, 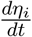 (red dotted lines), and distinguish the effects of natural selection, Δ_*ns,i*_ (black solid lines), changes in the environment, Δ_*ce,i*_ (blue dotted-dash line), and resource switching, Δ_*rs,i*_ (green dash lines). Note that because the environment includes both biotic and abiotic forces, the effect of environmental change on the average fitness of the population can be further decomposed into strain level effects.

## 4 Discussion

We provide a general framework to study the evolutionary dynamics of microbial communities and associated traits. Encouraged by the success of integrating population genetics into epidemiology [10, 18, 19], we first recast a classic resource-mediated population dynamic model in terms of evolutionary forces and then add traits to this framework. These models are particularly insightful if the dynamics of mean fitness and/or traits are of interest. This approach complements studies of long-term system behavior as it allows for the analysis of transient dynamics and reinterpretation of predictions derived from equilibrium analyses.

Rewriting population dynamic equations in terms of the proportion of microbial abundance in a given group uncovers relationships governing the evolutionary dynamics of the system explicitly in terms of the fitness of each group relative to the mean fitness of the entire population. Specifically, the dynamics of the mean fitness of the entire population are driven by three processes: natural selection, mutation, and changes in the environment. At equilibrium, these three forces balance. Given perturbations, different forces dominate depending on the nature of the perturbation. These forces are intertwined in that modulating one (e.g., introduction of novel species or changes in resource supply levels) lead to dynamics changes in the other forces.

We also provide an alternative form of the general model that reflects multi-level selection. We show that the strength of these three evolutionary forces may vary between subgroups (strains, species, etc). This approach helps to understand the impact of variable environmental conditions on community dynamics and compositions via feedback between ecological and evolutionary dynamics.

Our work is based on an extension of Tilman’s classic resource competition model [50]. It incorporates metabolic byproducts and numerous types of interactions (e.g., competition, mutualism, commensalism, amensalism). In our model, microbial interactions are indirect and mediated by molecules in the environment (e.g., nutrients, toxins), rather than direct and constant as assumed in generalized Lotka-Volterra and multivariate autoregressive models. These molecules include resources that are externally supplied as well as metabolic byproducts, and could easily be extended to include other molecules such as nutrients and toxins.

Different environmental conditions and resource levels will allow the growth of some microbes but not others, leading to the selection of certain microbial traits for survival. In population genetics, Price’s equation decouples the evolutionary change in frequency of a trait into two sources. First, the covariance between fitness and trait represents selection on the trait. Second, an in-tragroup expectation term represents trait evolution due to other factors besides selection (e.g., mutation) [38]. Here, we derive an equation that tracks the rate of change of the average value of any quantitative trait of interest, leading to a continuous-time version of Price’s equation (Eq. (4)). Because this form of Price’s equation is written in terms of mechanistic model parameters, it gives insight into which factors drive trait dynamics. For example, one would think intuitively that the group of microbes with the lowest mortality rate should be the winner in a competition. But recall in Example 1, the group of microbes with higher mortality rate (*S*_3_) won the competition (Figures 1b and 2b), because the Malthusian fitness also depends on per capita reproduction rate and resource availability.

In our simulations, we focus on a few canonical interaction types commonly encountered in microbial communities. We begin with the classical case where microbes compete for nutrients with no metabolic byproducts. We were able to verify the competitive exclusion principle (*n* groups cannot coexist on fewer than *n* resources at stationary state) [2, 31] when mutation is absent. With mutation, groups that would otherwise go extinct can maintain low abundance (Example 1).

Studies have shown that perturbations of microbiomes can disrupt the balance of microbial communities and the symbiotic relationship between the host and associated microbes. Functionally, these perturbations can result in diseases such as inflammatory bowel disease [47], obesity [37], and colon cancer [41]. An understanding of transient dynamics is required to predict transitions from healthy states to states associated with dysbioses. Our model captures the transient dynamics when the system is not at equilibrium, e.g., the temporal bloom of certain groups of microbes that eventually are out-competed by other groups (Example B1). Given sufficient information on parameter values, our model can be used to make predictions about these transient dynamics.

Incorporating metabolic byproducts into the model allows us to explore a broader range of community dynamics for small communities with basic interaction types. We show that more than one group of microbes can be supported by one externally supplied resource through interactions such as cross-feeding, and that environmental conditions can influence species dynamics and longterm equilibrium composition. If resources are supplied at sufficiently low constant rates, the community may collapse; as resource supply rates increase to a sufficiently high level, the community will reach an equilibrium state and stabilize. The time needed to reach equilibrium decreases as the resource supply rates increase and with overall higher microbial abundances. Butler and O’Dwyer also identified the role of external supply rate on community stability in their model of competition through exchange of resource [8]. On the other hand, time-varying resource supply rates lead to fluctuations in microbial community dynamics, as shown in Section 3.2–3.3. This framework allows us to explore how changes in the environment affect community dynamics, microbial interactions, and even coexistence, as normally microbes do not grow in a resource replete environment and therefore resources are not supplied at constant rates as in a chemostat.

We demonstrated that some traits may drive evolution directly, such as the net reproductive rate *γ = ϵα* and mortality rate *μ*. We can also track the dynamics of the mean value of traits that do not directly drive selection and predict the rate of evolution. For example, different groups of microbes may have different antibiotic tolerance levels, which is not the direct force driving selection in an antibiotic-free environment. Yet we can still track the mean value of antibiotic tolerance of the microbiome over time which could be useful to predict how this particular microbiome will withstand the introduction of antibiotics. Price’s equation provides an informative perspective on the relationship between trait dynamics and community dynamics, it also allows us to explore whether interventions such as modifying the microbial composition can promote health. A quantitative measure of traits associated with each microbial strain and knowledge of their population dynamics will allow us to predict the speed of evolution, and thus shed light on the link between microbial community composition and community function. There is an increasing interest in applications of trait-based approaches in microbial studies [30, 35], and a growing set of tools to evaluate and study microbial traits such as the Biolog Plates Technique [20, 46] and functional profiling with metagenomics and -omics data [17]. These tools offer great potential to identify microbial traits that are important to ecosystem functions and measure these functional traits.

Our work relies on several assumptions. One assumption is that the systems are well-mixed. That is, individual microbes have equal access to nutrients and toxins and interact with equal probability with all other microbes. We further assume that microbes in the same group have the same functional response and values of parameters such as reproduction rate, mortality rate, metabolic-byproduct excretion rate, and trait values. A natural extension of this framework would be to account for spatial and/or temporal heterogeneity. In our simulation studies, another assumption is that the growth rate function takes the form of a linear functional response. Toxins are assumed to increase mortality rather than decrease growth rate. Our model can be extended to include more realistic and complex functional responses, such as allowing for the saturation of resource utilization and different types of inhibition, incorporating more realistic metabolic networks such as co-limitation by multiple resources [27, 43].

While this model provides a robust conceptual framework, it will be challenging to connect it to data for particular systems as many parameters may be unknown. For example, the ability to metabolize complex compounds and the metabolic byproducts that each strain produces in a gut microbiome will typically be unknown. Additionally, not all bacteria can be grown in the laboratory with current techniques, and these unculturable bacteria may play critical role in maintaining the balance of ecosystems and health of their hosts. Yet the modeling framework presented in this paper highlights the key parameters that should be measured to assess potential intervention strategies. As more types of data become commonly collected in conjunction with microbial abundance data, trait databases could be constructed. Such tools could then lead to a more comprehensive understanding of the interactions governing the dynamics of microbial species and their surrounding environment, which is the key to building predictive models linking microbial community composition to functional response. Our framework can be adapted to microbial communities where manipulation to provide specific ecosystem services is desired, thereby facilitating the development of effective and precise microbiome-based interventions to promote health or other beneficial states.

## A Alternative Form and Multi-level Selection

Let *S_i_* denote the abundance of group *i* and *S_ik_* denote the abundance of subgroup *ik*, with 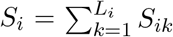. Model (1) can be rewritten as

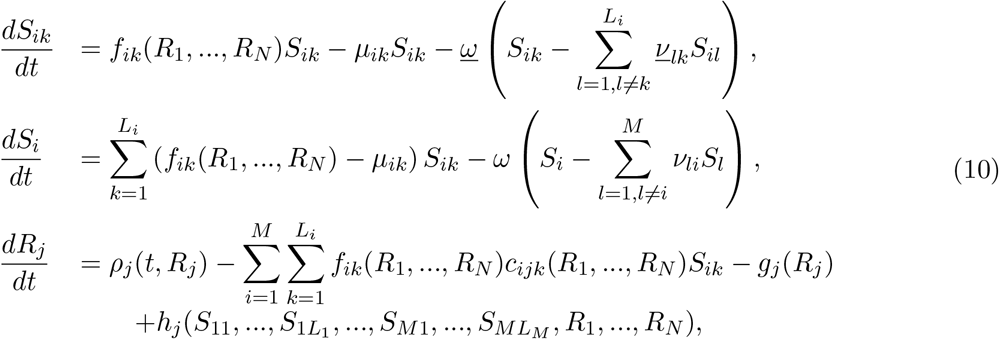

for *i* = 1,…, *M, j* = 1,…, *N, k* = 1,…, *L_i_*, where *L_i_* denotes the total number in subgroups of group *i*, 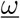 denotes the mutation rate of each subgroup, 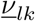 denotes the probability that *S_il_* mutate into *S_ik_, ω* denotes the mutation rate of each group, *ν_li_* denotes the probability that *S_l_* mutate into *S_i_*. Define 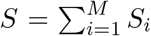 as the total population, *p_ik_* = *S_ik_/S_i_* as the frequency of subgroup *ik* within group *i, q_i_* = *S_i_/S* as the frequency of group *i*, we obtain

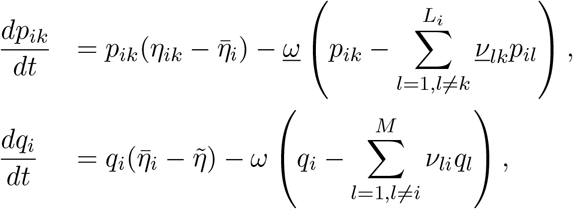

where

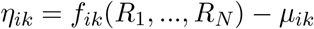

is the Malthusian fitness of 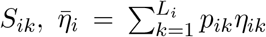 is the average Malthusian fitness of *S_i_*, and 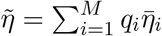 is the average Malthusian fitness of *S*. The temporal dynamics of 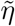 are now

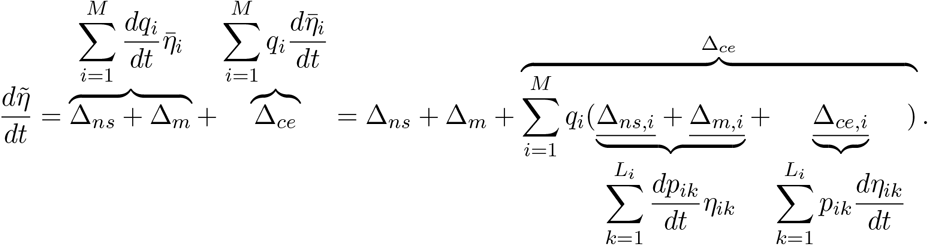

## B Competition without metabolic byproducts

### Example B1

In this example, two groups of microbes compete for a single resource, no mutation is allowed, that is, *N* = 2 and *ω* = 0 in (6), other parameter values and initial condition are listed in Table 4. At equilibrium, *S*_1_ has the lowest value of *R*\*, the numerical simulation confirms that *S*_1_ outcompete *S*_2_ in the end, as shown in Figures 7a. But the values of *R*\* does not provide information on the short term evolutionary dynamics of the competing groups of microbes before reaching the equilibrium.

**Figure 7:**
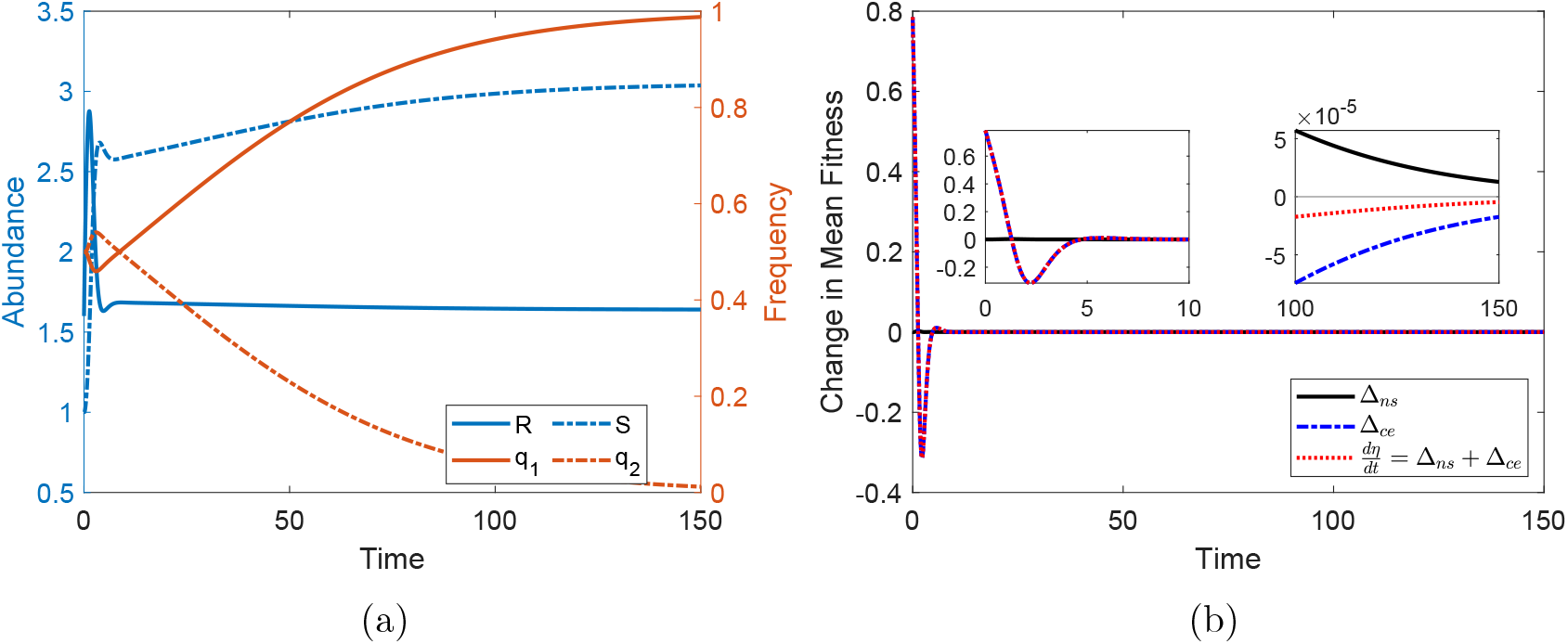
Classical resource competition. Numerical simulation of Example B1 shows the outcome of two groups of microbes compete for one resource, no mutation is allowed. (a) Abundance of resources and total microbial population (left axis) and frequencies of each group of microbes (right axis) over time. (b) Change in mean fitness 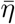 and effects of different drivers over time.

**Table 3:**
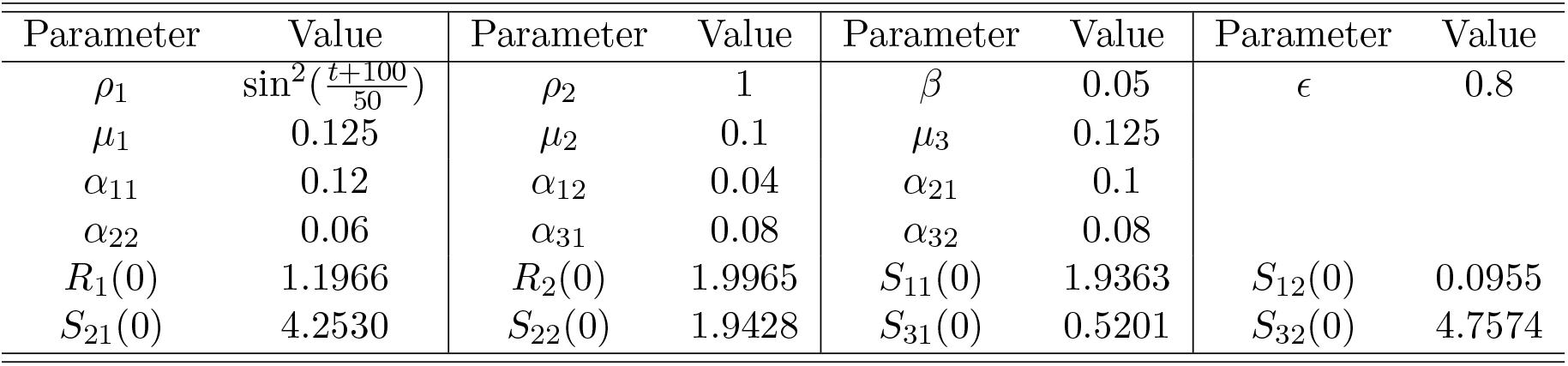
Parameter values and initial condition used in Figure 6.

**Table 4:**
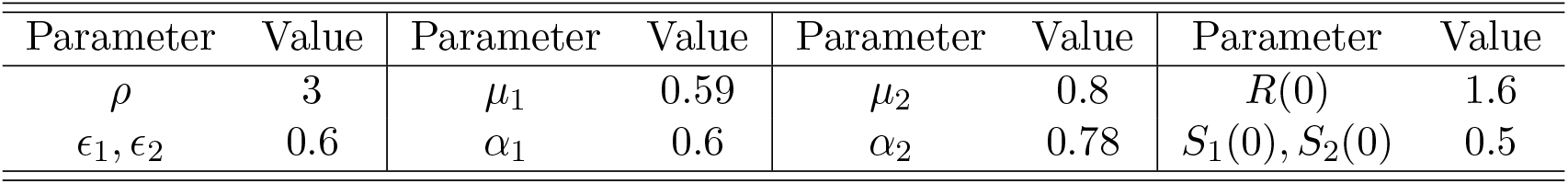
Parameter values and initial condition used in Figure 7.

Recall equation (7), natural selection favors in net reproductive rate favors *S*_2_ (*ϵ*_1_*α*_1_ > *ϵ*_2_*α*_2_), and in mortality rate favors *S*_1_ (*μ*_1_ < *μ*_2_). But high reproductive rate may be associated with high mortality rate, thus overall fitness may be low, as *S*_2_ in this example. Direct selection pulls net reproductive rate upwards with strength proportional to the resource abundance, *R*, while indirect selection drives net reproductive rate downwards with strength −1; for mortality rate, direct selection drives it downwards with strength −1, while indirect selection pulls it upwards with strength proportional to the resource abundance. Initially resource is abundant, there is little competition and both groups of microbes are able to grow at their maximal rate, the frequency of the group with the lowest fitness (*S*_1_) decreases, as indicated by the negative selection coefficient λ_1_ in equation (2). As the abundance of total population increases, the resource becomes scarce and microbes need to compete for resource. Initially, *S*_2_ reaches the highest frequency despite the fact that it has higher value of *R*^*^ than *S*_1_, because the resource abundance is relatively high, and the selection towards high *ϵα* is stronger than the selection towards low *μ*, in other words, *S*_2_ has a higher fitness than *S*_1_. But later on, the resource abundance decreases to be lower than *R*^*^ for *S*_2_, and the selection begins to favor *S*_1_ as it has higher fitness than *S*_2_ in this case, as shown in Figure 7a.

In short, the approach we introduced in Section 2 captures the transient dynamics which would be missed by the traditional equilibrium and near-equilibrium analysis, i.e., the temporal bloom of *S*_2_ which outcompete by *S*_1_ eventually in this case. This approach provides an explanation for the transient competitive advantage of certain group(s) of microbes, and allows us to predict the speed of evolution via equations (2)–(4), which can be valuable when making predictions about experimental manipulation of microbial communities with regard to health interventions.

## C Basic interaction types with metabolic byproducts

We now proceed to the extended case which include production of metabolic byproducts. As mentioned earlier, higher-order interactions also have impact on the functioning of microbial communities. Microbes can interfere or damage other microbes through production of toxic metabolites [21, 28, 40]. Some microbes may benefit from the metabolites secreted by other microbes [42]. For simplicity, we set mutation rate to be zero. Here we illustrate four basic types of interaction with metabolic byproducts, namely competition, mutualism, commensalism, and amensalism, as illustrated in Figure 8.

**Figure 8:**
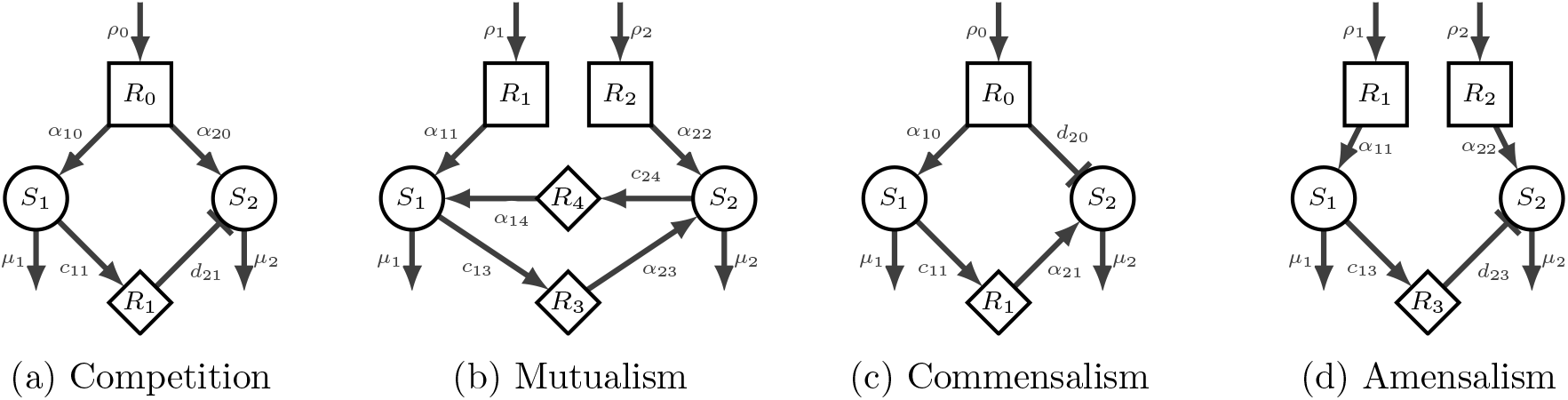
Illustration of the topological structure of the microbial communities discussed in Appendix C. Squared nodes denote externally supplied resources, circled nodes denote microbes, diamond nodes denote metabolic byproducts. Interactions are directed, positive interaction with an arrow head and negative interaction with a bar head.

### C.1 Competition with inhibition

In this example, 2 groups of microbes 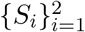 compete for 1 externally supplied resource, *S_i_* release toxic metabolic byproduct *R*_1_ which have negative impacts on *S*_2_. *R*_1_ can be a toxin that inhibit the growth of *S*_2_, or kill *S*_2_. For simplicity, we assume that the presence of *R*_1_ will increase the mortality rate of *S*_2_ by *d*_21_, as shown in Figure 8a. The model is as follows,

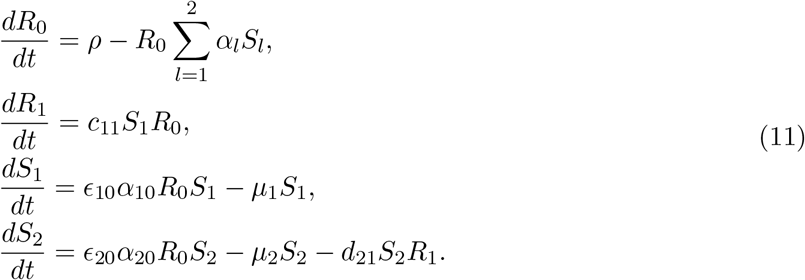

Key parameter values of the microbial groups are listed in Table 5. In the absence of the metabolic byproduct *R*_1_, *S*_2_ will outcompete *S*_1_ as it has lower value of *R*\*. When the influx rate of Ro is low, *ρ* = 0.01, the abundance of the metabolic byproduct *R*_1_ is very low, its negative effect on the growth of *S*_2_ is relatively low, *S*_2_ still has higher fitness than *S*_1_ and thus outcompete *S*_1_, as shown in Figure 9a. When we increase the influx rate of *R*_0_ to *ρ* = 0.1, initially *R*_0_ is abundant and both groups of microbes can grow at their maximum rate, the frequency of *S*_2_ increases as it has higher fitness. As the abundance of *S*_1_ increases, it produces more *R*_1_, and the strength of its negative impact on *S*_2_ increases which lead to a decrease in the fitness of *S*_2_. Once the accumulation of *R*_1_ reaches certain point such that the fitness of *S*_2_ is lower than that of *S*_1_, the frequency of *S*_1_ begins to increase and eventually outcompete *S*_2_, as shown in Figure 9b.

**Figure 9:**
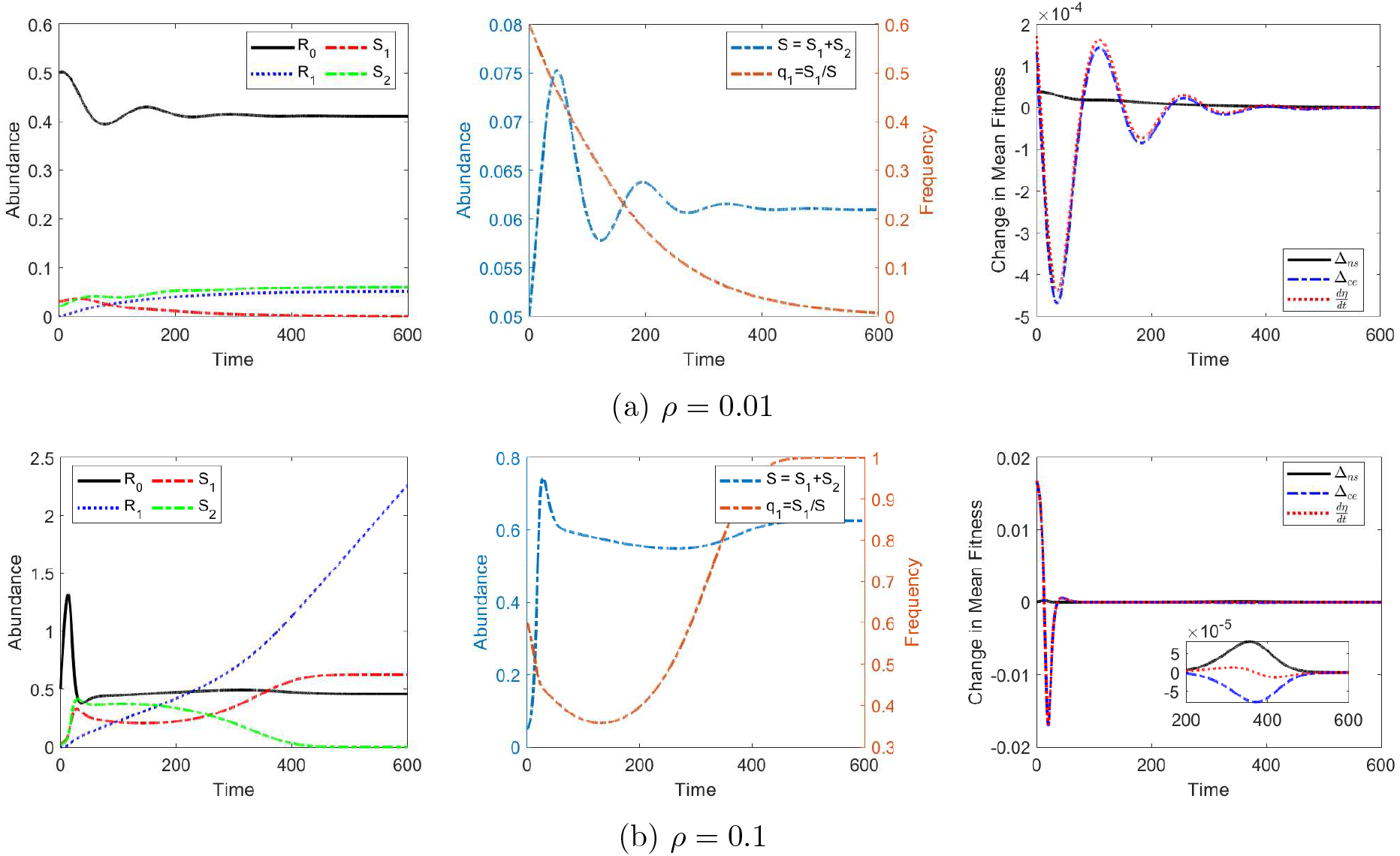
Numerical simulation of the competitive community, model (11). The values of the initial condition and all parameters except *ρ* are the same in both cases, as listed in Table 5. From (a) to (b), the resource supply rate, *ρ*, is increased by a factor of 10. Note that in the left two panels, solid lines denote resource, dotted lines denote metabolic byproduct, and dotted-dash lines denote microbial groups.

**Table 5:** Key parameter values and initial condition for the model (11) to produce solutions in Figure 9.

| Parameter | Value | Parameter | Value | Parameter | Value | Parameter | Value |
| --- | --- | --- | --- | --- | --- | --- | --- |
| $\mu_1, \mu_2$ | 0.08 | $\epsilon_1, \epsilon_2$ | 0.5 | $\alpha_{10}$ | 0.35 | $\alpha_{20}$ | 0.4 |
| $c$ | 0.02 | $d_{21}$ | 0.04 | $R_0(0)$ | 0.5 | $R_1(0)$ | 0 |
| $S_1(0)$ | 0.03 | $S_2(0)$ | 0.02 | | | | |

Note that if the growth rate function takes the form of the Holling Type II response, we can further consider the metabolic byproduct *R*_1_ as different types of inhibitors. If *R*_1_ is a competitive inhibitor to *S*_2_, then the growth rate of *S*_2_ in the presence of *R*_0_ and *R*_1_ is

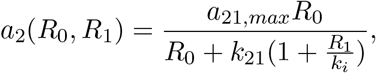

where *k_i_* is the inhibitory constant; if *R*_1_ is a non-competitive inhibitor to *S*_2_, then

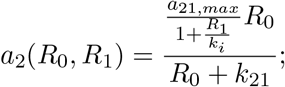

and if *R*_1_ is uncompetitive inhibitor to *S*_2_, then

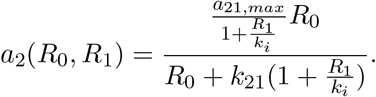

### C.2 Mutualism

This example consists 2 groups of microbes *S*_1_ and *S*_2_, each consumes one externally supplied resource *R*_1_ and *R*_2_, respectively. Each group of microbe releases one metabolic byproduct that will be utilized by the other microbe, as shown in Figure 8b. Assume that the effects from different resource on a focal microbe group are additive, we can derive the dynamic model of this community as

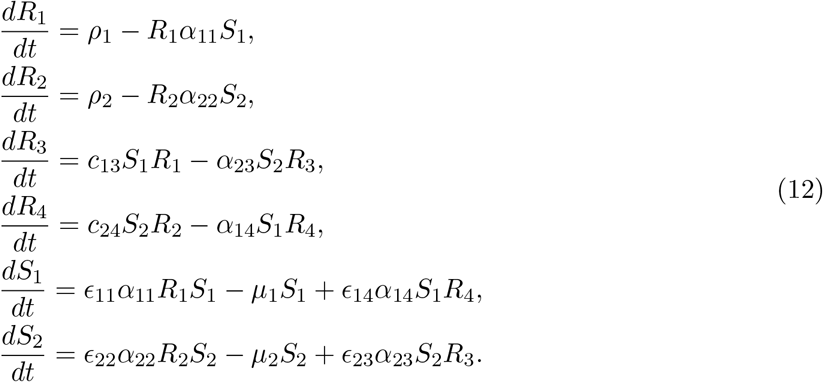

Here *S*_1_ and *S*_2_ will coexist because they don’t compete for resources. When the resource influx rates are low, *ρ*_1_ = *ρ*_2_ = 0.005, resources are scarce, the abundances of both groups of microbes decreases initially and then fluctuate, because they can utilize the limited amount of metabolic byproducts produced by the other group of microbes in addition to the externally supplied resources. But the overall abundance is low, as shown in Figure 10a. Note that even though change in the environment is the main driver of the change in mean fitness, with some small time ranges natural selection might be the dominant force. When we increase the resource influx rates to *ρ*_1_ = *ρ*_2_ = 0.05, initially the abundances of both groups of microbes decrease, as the resources are not abundant. But as resources are not as scarce as the first case, both groups of microbes are able to maintain relative higher abundance and produce more byproducts that can be utilized by the other group, the system reaches equilibrium state much quicker, as shown in Figure 10b.

**Figure 10:**
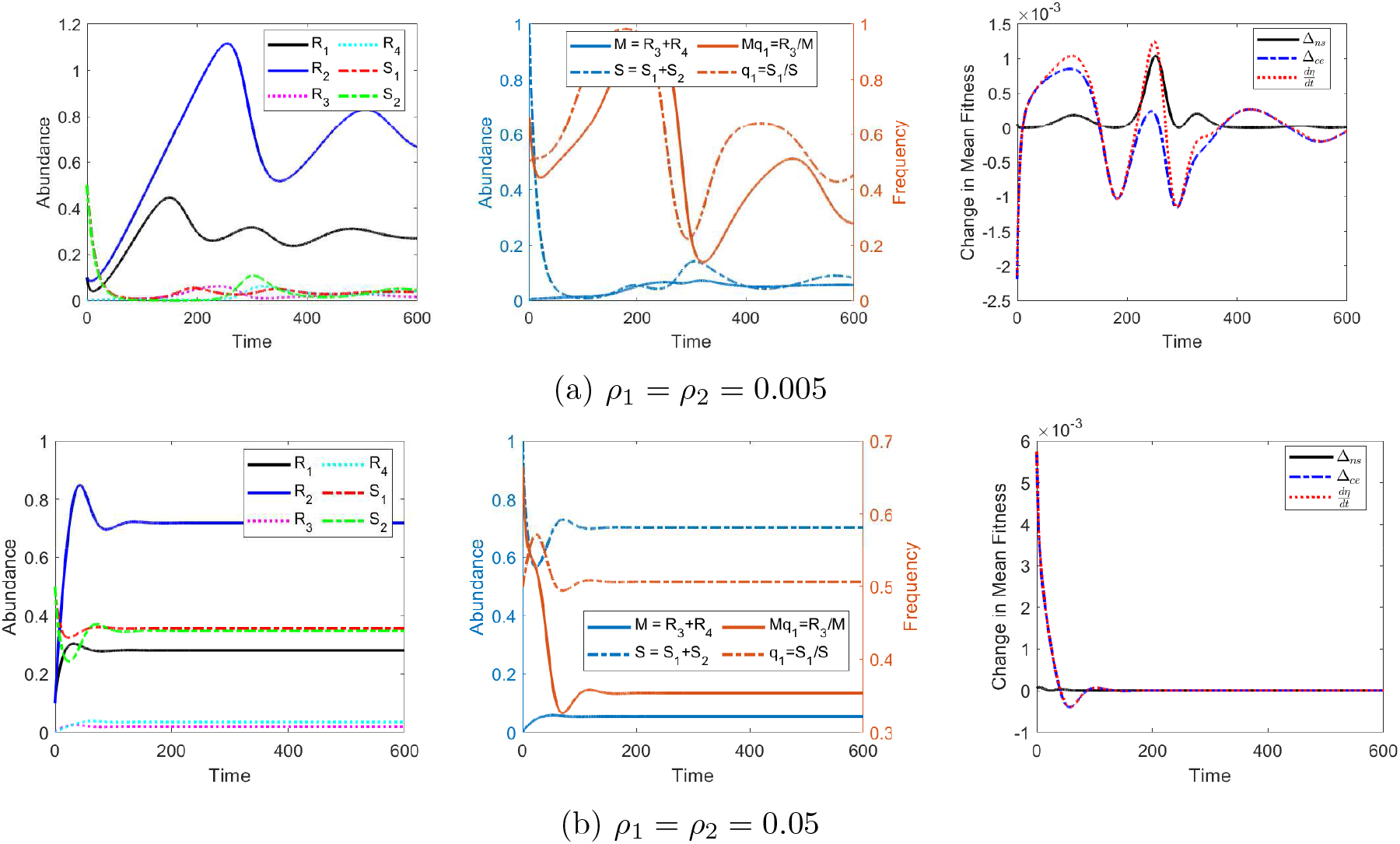
Numerical simulation of the mutualistic community, model (12). Key parameter values and initial conditions are listed in Table 6. From (a) to (b), the resource supply rates, *ρ*_1_, *ρ*_2_, are increased by a factor of 10. Note that in the left two panels, solid lines denote resource, dotted lines denote metabolic byproduct, and dotted-dash lines denote microbial groups.

**Table 6:** Parameter values and initial condition for the model (12) to produce solutions in Figure 10.

| Parameter | Value | Parameter | Value | Parameter | Value | Parameter | Value |
| --- | --- | --- | --- | --- | --- | --- | --- |
| $\mu_1, \mu_2$ | 0.08 | $\alpha_{11}$ | 0.5 | $\alpha_{23}$ | 0.6 | $R_0(0), R_1(0)$ | 0.1 |
| $c_{13}$ | 0.04 | $\alpha_{22}$ | 0.2 | $\epsilon_{11}, \epsilon_{22}$ | 0.5 | $R_3(0), R_4(0)$ | 0 |
| $c_{24}$ | 0.02 | $\alpha_{14}$ | 0.4 | $\epsilon_{14}, \epsilon_{23}$ | 0.7 | $S_1(0), S_2(0)$ | 0.5 |

### C.3 Commensalism

The fourth example consists 2 groups of microbes *S*_1_ and *S*_2_, and one externally supplied resource *R*_1_ which can be utilized by *S*_1_ and has negative impact on *S*_2_, respectively, and *S*_1_ releases metabolic byproduct *R*_1_ which can be utilized by *S*_2_, as shown in Figure 8c. Assume that the presence of *R*_0_ leads to increased mortality of *S*_2_ at the rate *d*_20_, we have

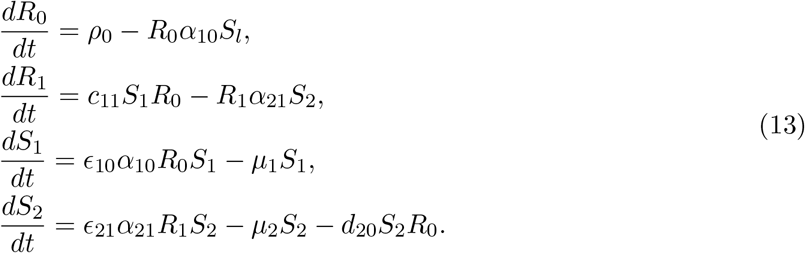

When *R*_1_ is absent or rare, *S*_2_ will go extinct. In Figure 11a, the influx rate of *R*_0_ is low, *ρ*_0_ = 0.01, and *R*_0_ is scarce initially (*R*_0_(0) = 0.04), abundances of both *S*_1_ and *S*_2_ decreases initially. During this phase, the amount of *R*_1_ produced by *S*_1_ is very low that it could not offset the negative impact of *R*_0_ on *S*_2_, so the abundance of *S*_2_ decreases quickly to zero. *S*_1_ dominates but with a low abundance, as shown in Figue 11a. When we increase the influx rate of *R*_0_ to *ρ*_0_ = 0.1, the external supplied resource *R*_0_ quickly build up, *S*_1_ can grow at its maximal rate shortly after the initial decrease phase, and produce enough *R*_1_ to support *S*_2_. In this case, *S*_1_ still dominate the population but both groups of microbes coexist and with higher abundances, as shown in Figure 11b.

**Figure 11:**
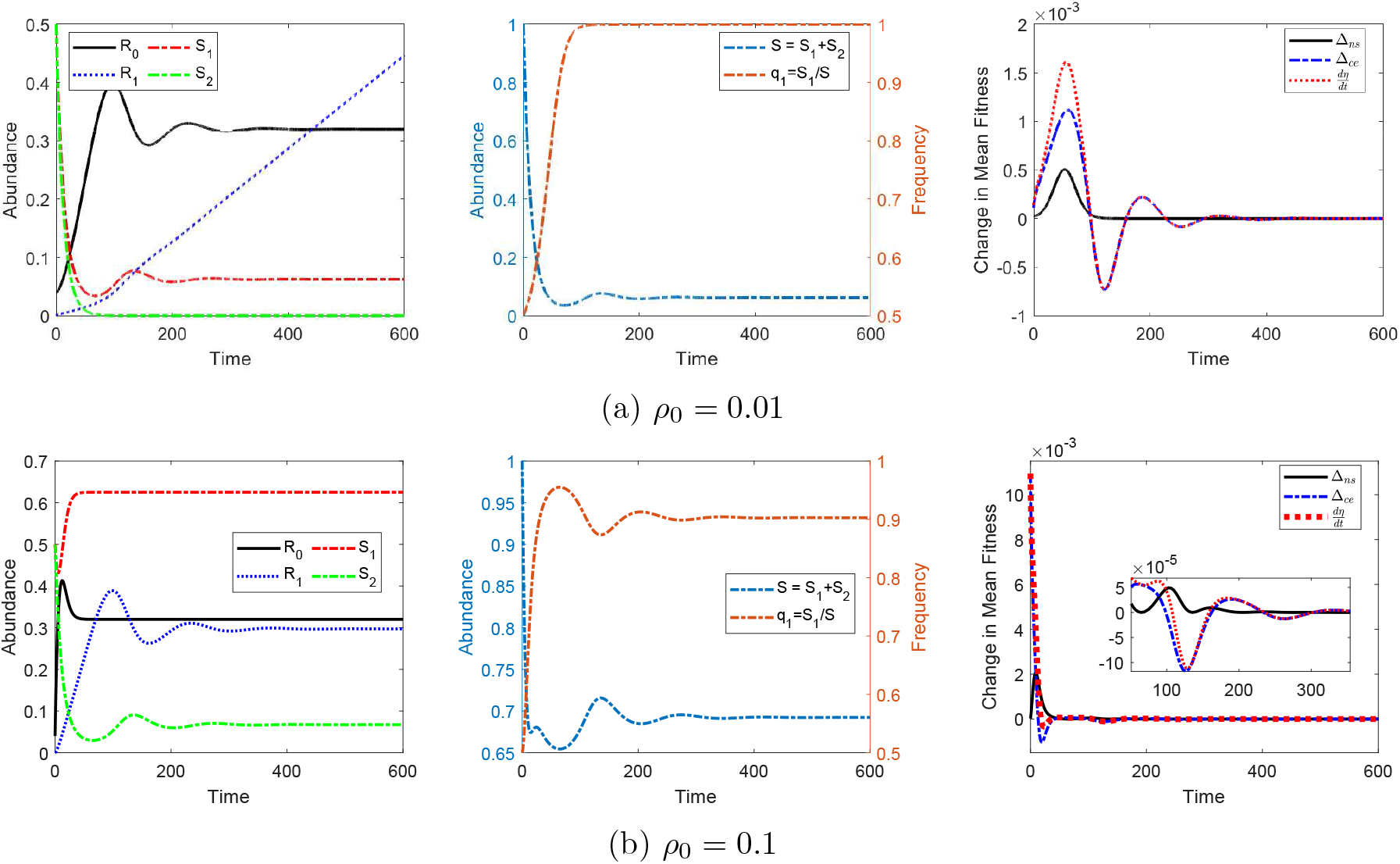
Numerical simulation of the commensalism community, model (13). Key parameter values and initial conditions are listed in Table 7. From (a) to (b), the resource supply rate, *ρ*_0_, is increased by a factor of 10. Note that in the left two panels, solid lines denote resource, dotted lines denote metabolic byproduct, and dotted-dash lines denote microbial groups.

**Table 7:** Parameter values and initial condition for the model (13) to produce solutions in Figure 11.

| Parameter | Value | Parameter | Value | Parameter | Value | Parameter | Value | Parameter | Value |
| --- | --- | --- | --- | --- | --- | --- | --- | --- | --- |
| $\mu_1, \mu_2$ | 0.08 | $\alpha_{10}$ | 0.5 | $\alpha_{21}$ | 0.4 | $d_{20}$ | 0.01 | $c_{11}$ | 0.04 |
| $\epsilon_{10}$ | 0.5 | $\epsilon_{21}$ | 0.7 | $R_0(0)$ | 0.04 | $R_1(0)$ | 0 | $S_1(0), S_2(0)$ | 0.5 |

### C.4 Amensalism

This example consists 2 groups of microbes *S*_1_ and *S*_2_, each consumes one externally supplied resource *R*_1_ and *R*_2_, respectively. *S*_1_ releases one metabolic byproduct *R*_3_ which have a negative impact on *S*_2_, as shown in Figure 8d.

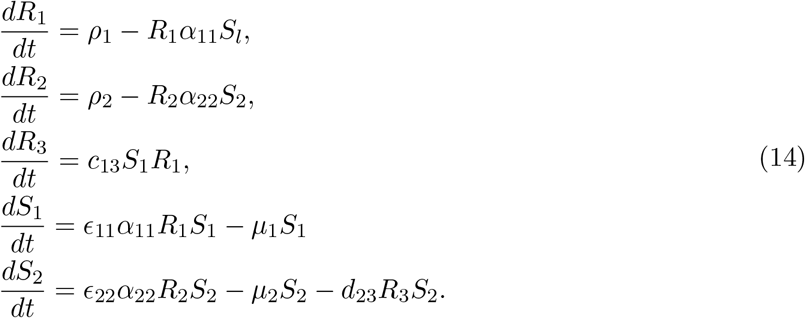

In the first case, *ρ*_1_ = *ρ*_2_ = 0.005, both resources are scarce. *S*_1_ maintain at a relatively low abundance, the amount of *S*_3_ accumulates in the environment is relatively low during the timespan in the simulation, and its negative impact on *S*_2_ is very small, both groups of microbes maintain at relatively low abundance, as shown in Figure 12a. When we increase the flux rate of *R*_1_ to *ρ*_1_ = 0.05, *S*_1_ is able to maintain a higher abundance and produce more *R*_3_, which drives the abundance of *S*_2_ to decrease quickly, as shown in Figure 12b.

**Figure 12:**
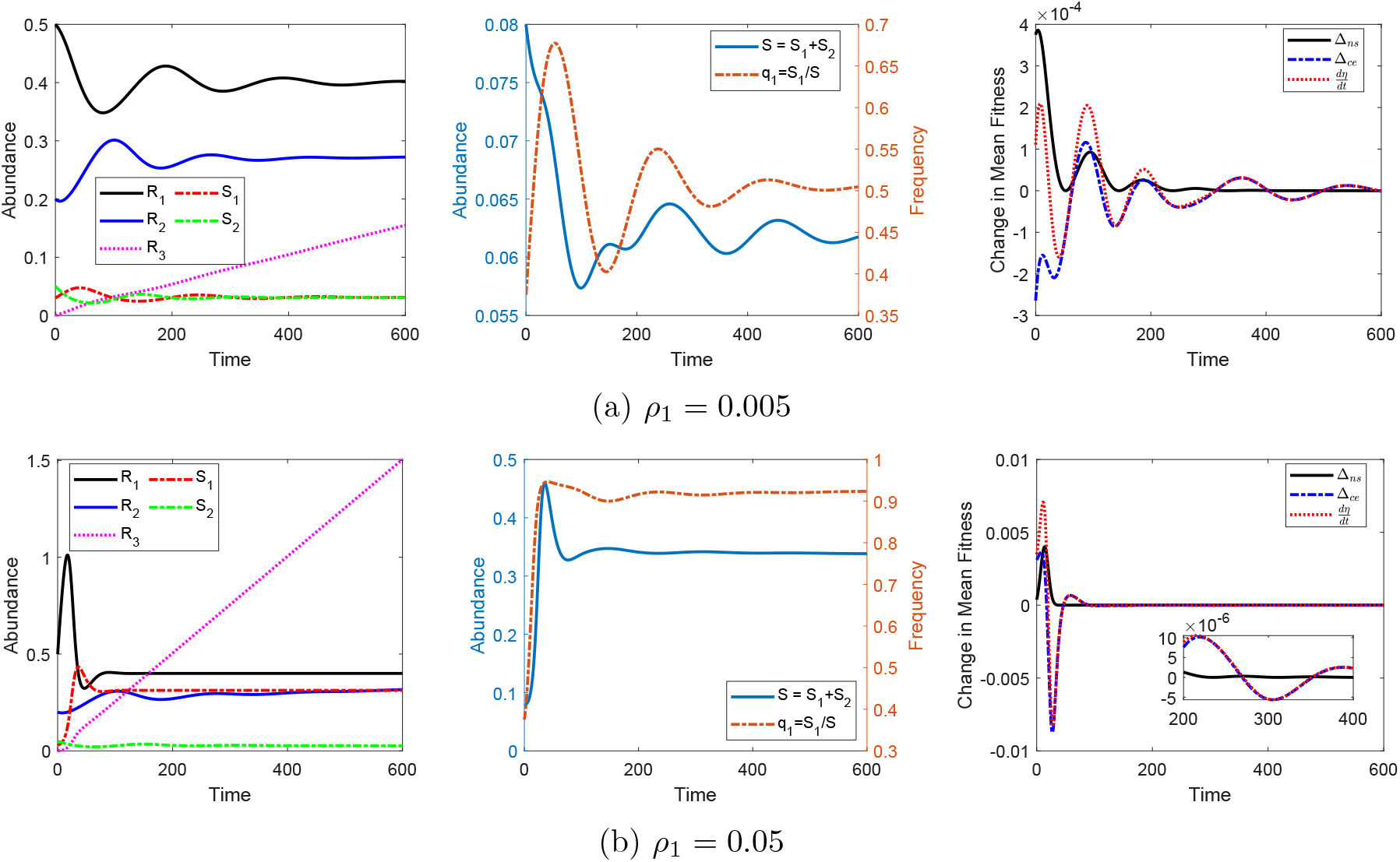
Numerical simulation of the amensalism community, model (14). Key parameter values and initial conditions are listed in Table 8. From (a) to (b), the resource supply rate, *ρ*_1_, is increased by a factor of 10. Note that in the left two panels, solid lines denote resource, dotted lines denote metabolic byproduct, and dotted-dash lines denote microbial groups.

**Table 8:**
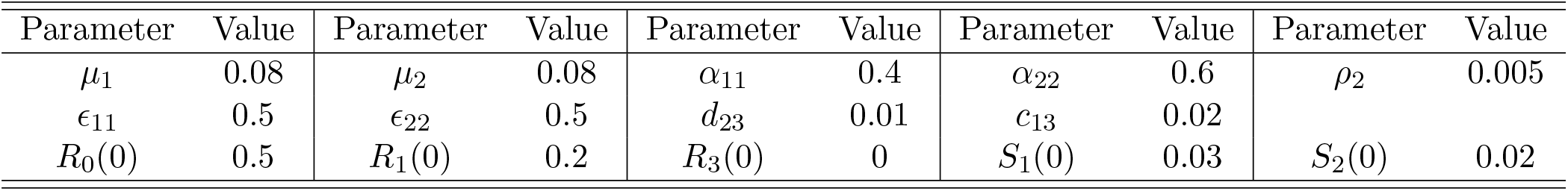
Parameter values and initial condition for the model (14) to produce solutions in Figure 12.

The communities in the examples shown in this paper are simple, as each one shows a different type of interaction. The method we presented can be used to explore the evolutionary and ecological dynamics of complex communities that incorporate multiple types of interactions.

## D Resource switching

Abundances and proportions of the community in Example 2.

**Figure 13:**
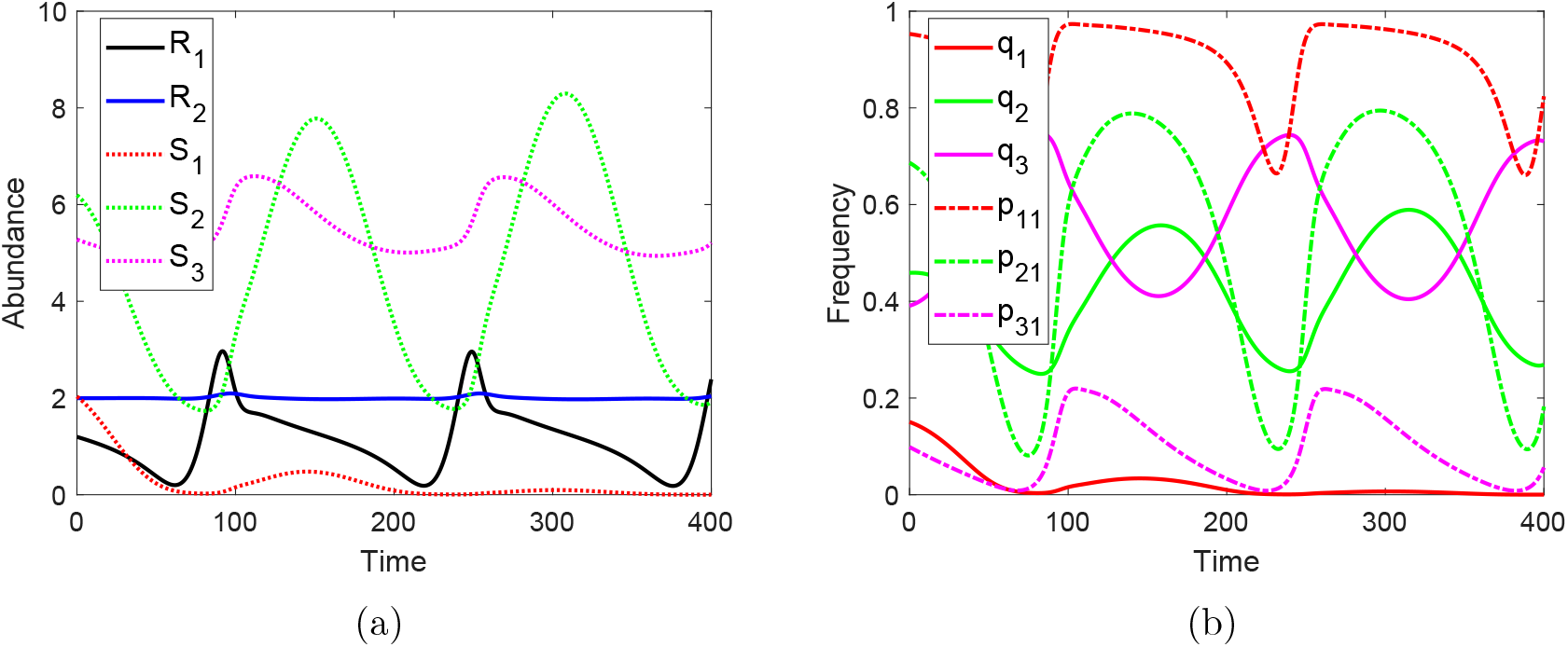
Numerical simulation of the community in Example 2. (a) Abundances of resources and microbes in different groups. (b) Proportions of microbes in different groups (solid lines) and subgroups that consume resource *R*_1_ (dotted-dash lines), the color of the lines identifies the group of microbes as in (a).

